# PerturbTrace: Evaluating Feedback Use by AI Co-Scientist Agents in Perturbation Discovery

**DOI:** 10.64898/2026.08.18.745260

**Authors:** Changjie Yu, Siyu Liu, Guanren Qiao, Mingrui Luo, Yujia Xiang, Zhiping Xu

## Abstract

Recent advances in AI co-scientists have brought LLM agents into closed-loop experimental design. However, whether these agents use feedback from earlier rounds to revise subsequent experimental decisions remains unclear. We address this question with PerturbTrace, which evaluates each round-to-round transition through Feedback-to-State, State-to-Action, and Action-to-Outcome. These stages assess whether feedback is reflected in the agent’s rationale and perturbation-selection strategy, whether the stated strategy guides the next perturbation batch, and whether that batch yields more hits than expected under random sampling. We evaluate four LLM agents on 17 screen-derived tasks and compare them with random selection, active learning, and LLM-guided Bayesian optimization baselines. Each agent outperforms the strongest non-agent method on at least 15 of the 17 tasks, yet controlled evaluations across six tasks show no consistent advantage from true feedback over random or no feedback. Among 576 transitions under true or random feedback, only 43 (7.5%) complete the full Feedback–State– Action–Outcome sequence, including 25 under random feedback. These findings show that high final recall does not necessarily indicate effective feedback use. They also highlight the need to evaluate closed-loop scientific agents by both their discovery performance and whether feedback changes their subsequent decisions.

## 1 Introduction

Scientific discovery is iterative, progressing from observation and hypothesis generation to experimental design, execution, and analysis. AI co-scientists combine LLMs with scientific knowledge, computational tools, and experimental platforms to support this process. In Lab-in-the-Loop workflows, results from one round are returned before the next round begins (Ghareeb et al. 2026; Wainrib et al. 2026). Later experimental choices can therefore respond to evidence that was unavailable when the initial plan was formed. Recent systems in biology and chemistry increasingly extend one-shot hypothesis generation into such closed-loop workflows (Boiko et al. 2023; Gottweis et al. 2026; Ghareeb et al. 2026; Huang et al. 2026). Final discovery performance directly measures fixed-budget utility, but it cannot determine whether new evidence informed later decisions. Similar performance may instead arise from prior knowledge, tool use, or a fixed selection strategy. Evaluating closed-loop AI co-scientists therefore requires separating discovery competence from feedback-use capability.

Perturbation discovery is a closed-loop process for identifying genetic or chemical interventions that produce a desired biological effect. Candidate perturbations are tested in sequential batches, and results from each round can guide the next selection. We refer to effective perturbations as hits and measure discovery performance by the hits identified within a fixed experimental budget. Active-learning and Bayesian-optimization methods explicitly update a model or acquisition rule from observed outcomes (Mehrjou et al. 2022; Lyle et al. 2023; Huang et al. 2024; Rubbi et al. 2026). LLM agents can use biological knowledge, analyze experimental data, and express selection rationales in natural language (Roohani et al. 2025; Hao et al. 2025). The evaluated interfaces record the next written rationale and accepted batch but expose no directly verifiable internal decision state. We therefore treat the rationale as an externalized decision state *S_t+1_* for evaluation.

Existing evaluations address only parts of this problem. DiscoveryBench and ScienceAgentBench assess data-driven scientific workflows (Majumder et al. 2025; Chen et al. 2025). Studies of whether feedback improves the performance of AI co-scientists have reached different conclusions. Gupta, Hartford, and Liu (2025) found that randomly permuting outcomes did not alter agent performance. In contrast, Wainrib et al. (2026) reported gains from candidate-level feedback under a different model and randomization protocol. Feedback sensitivity is therefore an empirical question that depends on the evaluated setting. Controlled endpoint contrasts can associate feedback conditions with final outcomes, but they do not reveal how delivered feedback appears in later decision artifacts. A unified evaluation must jointly record endpoint discovery, feedback-condition contrasts, and an observable trace from feedback to decision state, accepted action, and subsequent outcome.

We introduce **PerturbTrace, a framework for transition-level evaluation of feedback use in closed-loop perturbation discovery**. We also introduce PerturbTraceBench, a benchmark containing 17 screen-derived tasks with task-specific targets, candidate spaces, and hidden hit sets. PerturbTrace decouples task execution, the evaluated agent, oracle feedback, scoring, and trace collection. Its three-stage evaluation follows each round-to-round transition from Feedback-to-State through State-to-Action and Action-to-Outcome.

**Our experimental analysis showed that:**

1. **LLM agents identify more effective perturbations than sequential baselines.** Each evaluated agent outperformed the strongest non-agent method on at least 15 of 17 tasks. At least one agent achieved the best performance on every task. However, the best-performing agent varies across tasks, and stronger endpoint performance does not imply more effective feedback use.
2. **True feedback does not reliably improve final recall.** Across the six-task controlled evaluation, the contrasts varied across agents and tasks. No agent showed a consistent gain over both random and no feedback, and apparent gains were sensitive to individual tasks.
3. **Feedback-supported state updates rarely guide subsequent actions.** Only 43 of 576 feedback-bearing transitions completed the full Feedback–State–Action–Outcome sequence, including 25 under random feedback. State updates were substantially more common than implementation in the next batch. Thus, endpoint success or a coherent response alone did not establish feedback-guided adaptation.

**Our contributions include:**

1. **We introduce PerturbTrace, a Feedback–State–Action–Outcome framework for transition-level evaluation of feedback use in closed-loop scientific decision-making.** The framework distinguishes whether feedback supports a state update, whether that update guides the next action, and whether the action improves the subsequent outcome.
2. **We apply this framework to evaluate feedback use by LLM-based scientific agents in perturbation discovery.** True-feedback, random-feedback, and no-feedback conditions separate endpoint performance from observable feedback use and identify where the feedback-to-outcome sequence stops.
3. **We develop the PerturbTrace Harness and PerturbTraceBench for modular, reproducible evaluation.** The combined infrastructure supports one-command, end-to-end execution and extends to closed-loop scientific workflows in which experimental evidence guides subsequent decisions.

These results motivate evaluating closed-loop agents by both their discovery performance and the influence of experimental evidence on subsequent decisions. Figure 1 summarizes (A) the finite-budget perturbation-discovery loop, (B) the PerturbTraceBench task interface, and (C) the transition-level PerturbTrace audit.

**Figure 1:**
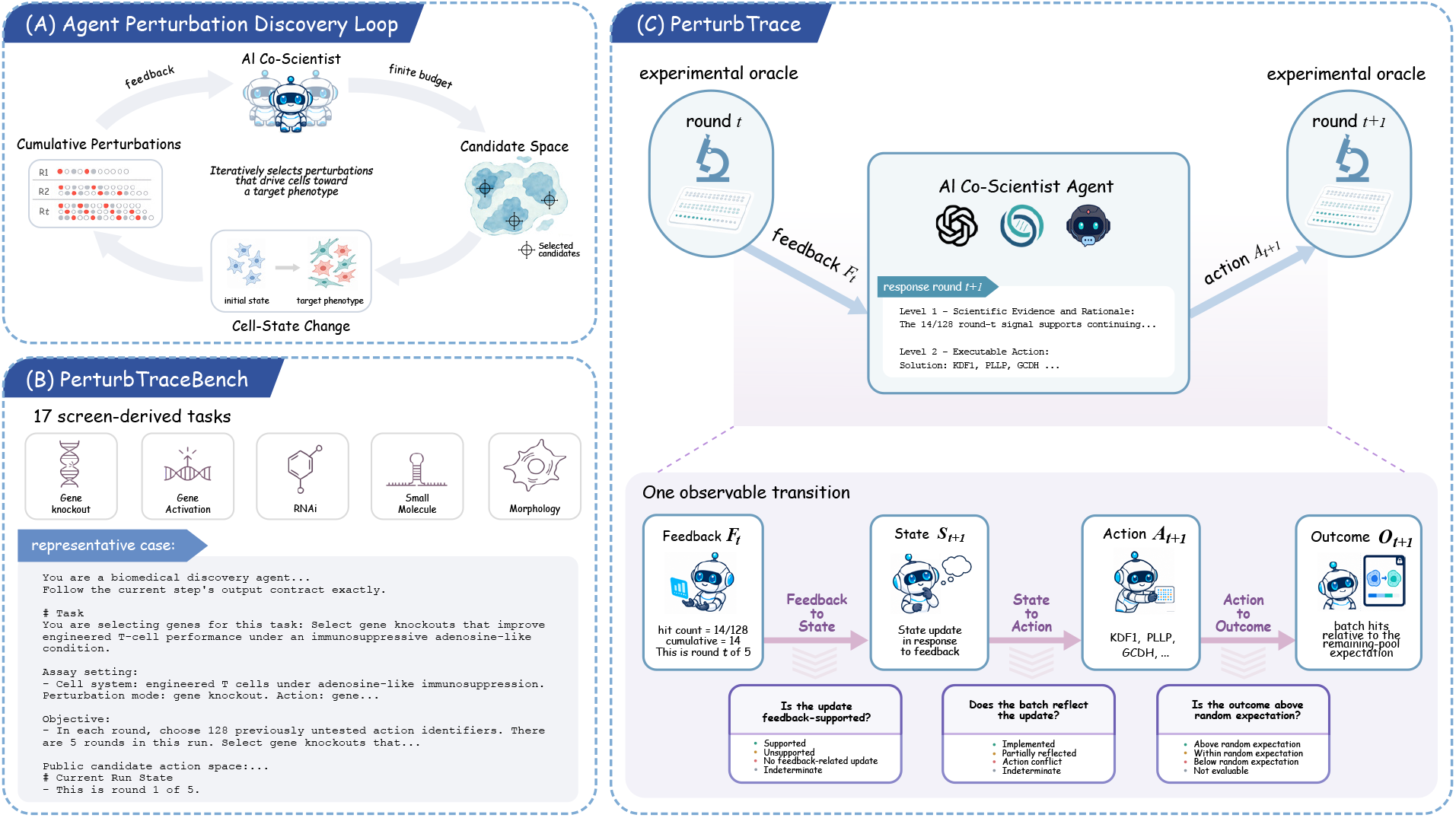
Overview of PerturbTraceBench and PerturbTrace. (A) Under a finite budget, an AI co-scientist iteratively selects perturbation batches from a candidate space and receives feedback. (B) PerturbTraceBench contains 17 screen-derived tasks spanning gene knockout, gene activation, RNA interference, small molecules, and morphology. The representative card shows the agent-visible task description. (C) PerturbTrace standardizes the interaction between an experimental oracle and an AI co-scientist. For each adjacent-round transition, it records the delivered feedback *F_t_*, the next rationale as decision state *S_t+1_*, the accepted batch *A_t+1_*, and evaluator-side outcome *O_t+1_*. It evaluates whether feedback supports the stated decision, whether the accepted batch reflects that decision, and whether the batch yields more hits than expected under random sampling.

## 2 Related Work

### Sequential perturbation discovery

Active learning and Bayesian optimization formalize experimental design as sequential selection under a limited budget. GeneDisco standardizes batch acquisition on drug-discovery screens, whereas DiscoBAX seeks effective perturbations across diverse mechanisms (Mehrjou et al. 2022; Lyle et al. 2023). IterPert selects informative perturbations to improve response prediction over short experimental horizons. Many Needles instead formulates threshold-based hit discovery as the acquisition objective (Huang et al. 2024; Rubbi et al. 2026). These methods incorporate observed outcomes through statistical models and acquisition rules. In contrast, PerturBench and Systema evaluate perturbation-response prediction rather than the sequential selection of experiments (Wu et al. 2025; Vinas Torne et al. 2026). PerturbTrace asks whether the next decision of an LLM agent reflects delivered feedback and is implemented in the accepted experiment.

### AI agents for scientific discovery

Tool-augmented LLM systems now support chemical planning, biomedical hypothesis generation, workflow execution, and laboratory-guided iteration (Boiko et al. 2023; Bran et al. 2024; Gottweis et al. 2026; Ghareeb et al. 2026; Huang et al. 2026; Qu et al. 2026). BioDiscoveryAgent and PerTurboAgent are closest to our setting. Both use LLM reasoning and biological resources to select perturbations over multiple experimental rounds (Roohani et al. 2025; Hao et al. 2025). These studies show that agents can produce useful hypotheses and experimental choices. PerturbTrace examines whether delivered feedback is reflected in the subsequent decision of an agent and the accepted perturbation batch.

### Scientific-agent evaluation and feedback use

DiscoveryBench evaluates multi-step hypothesis discovery, ScienceAgentBench evaluates executable scientific workflows, and WetBench provides a simulated environment for experimental planning (Majumder et al. 2025; Chen et al. 2025; Brown, Bracha, and Boyden 2025). More directly, Gupta, Hartford, and Liu (2025) randomized candidate outcomes and found little effect on LLM-based experimental design. Wainrib et al. (2026) later reported positive effects under a different model, candidate space, feedback channel, and randomization protocol. Together, these findings indicate that feedback sensitivity depends on the agent and experimental setting. PerturbTrace complements this work by evaluating screen-derived tasks across agent configurations with random- and no-feedback controls. It reports endpoint discovery, feedback-condition contrasts, and a transition trace. Feedback-to-State and State-to-Action provide behavioral evidence of feedback use, whereas Action-to-Outcome evaluates the immediate productivity of the response.

## 3 Methodology

### 3.1 Evaluating Feedback Use in Closed-Loop Perturbation Discovery

We define a multi-round perturbation-discovery task as 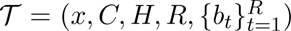, where *x* is the scientific target, *C* is the candidate space, *H ⊂ C* is the screen-derived hidden hit set, *R* is the number of rounds, and *b_t_* is the action budget at round *t*. At each round, an agent selects a batch *A_t_*from the untested candidates; the assigned feedback condition determines the information returned. Endpoint metrics quantify final hit discovery but cannot determine whether feedback changed later decisions. We distinguish discovery performance, measured by final hit discovery, from feedback use, assessed through observable links among experimental outcomes, subsequent rationales, and perturbation choices. High discovery performance therefore does not necessarily imply feedback use.

For each completed run, we measure discovery performance using final top-effect recall,

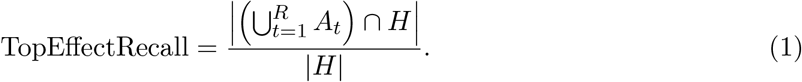

Here, *A_t_* is the perturbation batch submitted in round *t*, and *H* is the hidden hit set. The numerator counts the unique hits discovered by the final round, whereas the denominator is the total number of hidden hits in the task.

We evaluate feedback use through transitions between adjacent rounds:

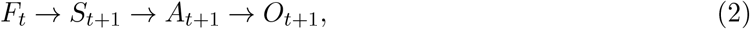

where *F_t_* is the delivered feedback and *S_t+1_* is the externalized decision state expressed in the next scientific rationale. The accepted perturbation batch is *A_t+1_*, and its experimental outcome is *O_t_*_+1_.

For each transition from round *t* to round *t* + 1, PerturbTrace records the feedback after round *t*, the next rationale, the accepted batch, and the outcome of that batch. We then ask whether the rationale relates the feedback to the next search direction and whether the accepted batch follows that direction. When both relations hold, the record provides observable evidence that feedback was translated into a stated decision and action. The *A_t+1_ O_t_*_+1_ step asks whether the batch discovers more hits than an equally sized random batch from the remaining candidate pool. This comparison measures the immediate productivity of the revised selection under the fixed-budget hit-discovery objective.

Interpreting the first two stages as evidence of feedback use requires the rationale to represent the decision associated with the final accepted batch. It also requires the annotation scheme to capture the stated search direction and its implementation. PerturbTrace therefore evaluates an externalized behavioral proxy under these assumptions rather than latent internal belief revision.

### 3.2 Overview of PerturbTrace Framework

PerturbTraceBench contains 17 tasks derived from real perturbation screens. The tasks span gene knockout, gene activation, RNA interference, small-molecule treatment, and morphology-based screening. Each task specifies a scientific target, a candidate perturbation space, and a screen-derived hit set. As illustrated in Figure 1(B), agents receive the task description, budget, and candidate identifiers, but not the scores, hit labels, ranks or thresholds of untested candidates.

Several tasks use resources that profile multiple cell lines. Each such task fixes one cell context rather than pooling measurements across cell lines. For GDSC2, DepMap/Achilles, and Project DRIVE, we selected the context with the greatest non-null candidate coverage in the source release. These tasks include GDSC2 drug response in HT-29 (SIDM00136) (Iorio et al. 2016), DepMap/Achilles CRISPR dependency in OVCAR-8 (ACH-000696) (Tsherniak et al. 2017), and Project DRIVE RNAi dependency in SF-295 (McDonald et al. 2017). JUMP Cell Painting contributes CRISPR morphology profiles from U2OS cells (Chandrasekaran et al. 2023), and the Wang CRISPR survival screen uses KBM7 cells (Wang et al. 2014).

PerturbTrace is a one-command, end-to-end evaluation framework with an execution harness and a three-stage trace evaluation. PerturbTraceBench provides the tasks. The harness standardizes task loading, multi-round interaction, oracle queries, feedback control, response validation, scoring, and trace collection. A common solver interface supports the evaluation of general-purpose agents under the same task, oracle, feedback rule, and scoring protocol.

True and random feedback provide the latest-round hit count and the cumulative hit count. True feedback computes both counts from the actual action-outcome mapping. Random feedback applies a fixed, run-seeded global permutation to this mapping and computes the same counts from the permuted mapping. The no-feedback condition provides no experimental outcome signal.

We conducted feedback-condition experiments on a prespecified six-task panel spanning variation in candidate-space size, hit sparsity, and perturbation modality. All panel tasks use gene-level actions, five rounds, 128 actions per round, and a 640-action budget. Candidate-space coverage equals the budget divided by the candidate count, whereas hit prevalence equals the hit-set size divided by the candidate count. The panel contains two tasks from each of three task-property strata. The budget-easy stratum contains tasks with high candidate-space coverage. The first is CRISPR selection in human cells (7,114 candidates, 321 hits, and 9.00% coverage). The second is Project DRIVE RNAi dependency (7,975 candidates, 399 hits, and 8.03% coverage). The standard genome-scale stratum includes HUPT3 CAR-T resistance (20,079 candidates, 884 hits, and 4.40% prevalence). It also includes NK-cell exposure CRISPR (20,617 candidates, 1,031 hits, and 5.00% prevalence). The feedback-sparse stratum contains CRISPRa exhaustion (18,797 candidates, 149 hits, and 0.79% prevalence). It also contains phagocytosis CRISPR (20,398 candidates, 260 hits, and 1.27% prevalence). The panel spans CRISPR knockout, CRISPR activation, and RNA interference under a common budget.

### 3.3 Agent configurations and Selection

We evaluated four agent configurations under a common GPT-5.5 backend with xhigh reasoning effort. BioDiscoveryAgent represents a biomedical discovery workflow with draft critique, whereas Biomni represents a biomedical agent with tool retrieval. Codex uses a general-purpose agent configuration with a task-neutral execution instruction. Codex+strategy uses the same Codex backend, reasoning setting, tool access, and task interface, with an additional restricted perturbation-selection strategy. Together, these configurations cover biomedical discovery workflows, tool retrieval, general-purpose agent operation, and an explicit decision procedure. All configurations use the same harness, task inputs, budgets, feedback policies, response validation, and scoring protocol.

### 3.4 Three-Stage Trace Evaluation

The three-stage evaluation applies the trace shown in Figure 1(C) to each adjacent-round transition. It asks whether feedback is reflected in the stated decision, whether that decision is carried into the accepted batch, and whether the batch yields an immediately productive discovery outcome.

#### Feedback-to-State: Does feedback enter the next decision?

We compare the delivered feedback with the scientific rationale in the following round. The rationale must interpret the feedback correctly and state how it supports maintaining or revising the current search direction. Feedback use does not require a directional change. Retaining a strategy can also be feedback-driven when the rationale provides an explicit link. If a clear claim lacks support from the delivered feedback, the transition is marked as unsupported. If the text is insufficient to determine whether feedback informed the rationale, the transition is marked as indeterminate.

#### State-to-Action: Do the next perturbation choices follow the stated decision?

We extract the next search direction from the scientific rationale and assign candidates in the accepted batch from the same round to task-specific biological categories. We then compare the stated direction with the batch composition. For example, if the rationale proposes expanding the search within a biological category, the next batch should contain more candidates from that category. This stage distinguishes a written response to feedback from a corresponding change in perturbation choices.

#### Action-to-Outcome: Does the next batch discover more hits than expected under random selection?

We compare the hit rate of the next batch with that of the remaining candidate pool. Let *N_t_* and *H_t_* denote the numbers of remaining candidates and hits before round *t* + 1. Let *b_t_*_+1_ and *h_t_*_+1_ denote the batch size and observed hit count. The relative hit yield is

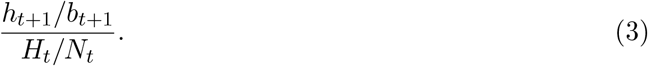

We use *X_t+1_*~ Hypergeom(*N_t_, H_t_, b_t_*_+1_) to classify the observed hit count as above, within, or below random expectation. An outcome is above random expectation when the upper-tail probability is at most 0.025. It is below random expectation when the lower-tail probability is at most 0.025. If no hidden hits remain, the relative hit yield is undefined, and Action-to-Outcome is marked as not evaluable.

Feedback-to-State and State-to-Action jointly provide observable evidence of feedback use. They assess whether feedback enters the next scientific rationale and is reflected in the next perturbation batch. Action-to-Outcome asks whether these choices also improve immediate discovery. We define productive full-chain completion as a transition that satisfies the first two stages and produces a hit count above random expectation. The outcome stage is not necessary for feedback use, and we therefore report all three stages separately. Under no feedback, Feedback-to-State and full-chain completion are not applicable.

We prespecified the observable evidence required at each stage and the handling of insufficient evidence. We evaluated a subset of 288 transitions using two independent judge passes. This subset comprised 144 development and 144 confirmation transitions. We revised decision rules only on the development set. The rules had to meet a prespecified agreement threshold on the confirmation set before application to the remaining transitions. The supplementary material provides the detailed decision rules, unclear cases, and agreement calculations.

PerturbTrace reports endpoint discovery and transition-level feedback-use results separately. Endpoint results describe final hit discovery. Transition-level results show whether feedback was reflected in the next state and action and whether the resulting action improved immediate discovery. We ran each feedback condition independently while holding the task, agent configuration, and budget fixed.

## 4 Experiments

Our evaluation addresses three questions. We first compare the overall perturbation discovery performance of LLM agents with methods that explicitly update a model across rounds. We then compare final performance under true feedback with performance under random or no feedback. Finally, we examine whether feedback informs the next perturbation choices and whether those choices improve the following-round outcome.

The overall evaluation covers 17 perturbation discovery tasks. It compares four LLM agents with Random, two active-learning methods, and an LLM-guided Bayesian-optimization method. All methods use three seeds and a fixed five-round budget. On six tasks, we additionally evaluate the four agents under true, random, and no feedback, yielding 216 runs. We analyze 576 adjacent-round transitions from the true- and random-feedback conditions to assess responses to feedback.

### 4.1 LLM Agents Identify More Effective Perturbations than Sequential Baselines

Figure 2 compares all methods across 17 perturbation-discovery tasks using hit recall after five rounds. Under the evaluated interfaces, each LLM agent outperformed the strongest non-agent method on at least 15 of 17 tasks. Biomni and Codex did so on 16 tasks, whereas BioDiscoveryAgent and Codex+strategy did so on 15 tasks. On every task, at least one agent outperformed all non-agent methods. The median task-level improvement was 9.3 percentage points in recall.

**Figure 2:**
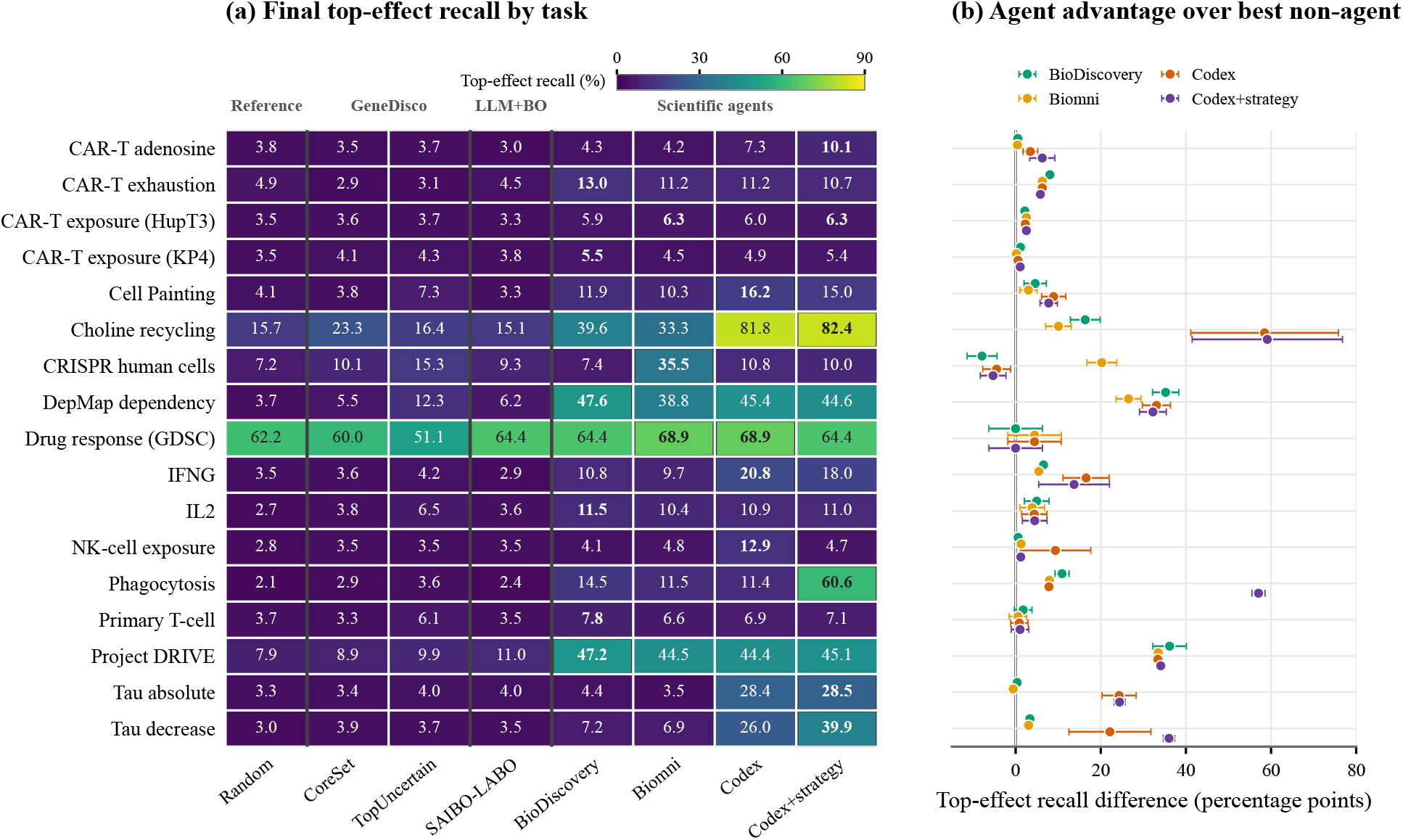
Task-wise discovery performance. Each task is evaluated in one fixed experimental context. (a) Mean final top-effect recall over three seeds on 17 tasks. Black outlines mark the highest mean per task. (b) Recall difference between each agent and the strongest non-agent comparator on the same task, selected from Random, the two GeneDisco methods, and SAIBO-LABO. Points compare three-seed means; whiskers show one standard error. Positive values favor the agent. CoreSet and TopUncertain denote the two GeneDisco acquisition strategies; BioDiscovery denotes BioDiscoveryAgent.

No agent performs best across all tasks. After adjustment for ties, BioDiscoveryAgent achieved the most task-level wins (6.0), followed by Codex+strategy (5.5), Codex (3.5), and Biomni (2.0). Codex+strategy nevertheless achieved the highest macro-averaged recall across the 17 tasks (27.3%). BioDiscoveryAgent won more tasks despite a lower macro recall (18.1%). Thus, large gains from Codex+strategy on a small subset of tasks influenced the aggregate ranking. Among classical sequential decision-making methods, SAIBO-LABO achieved 8.7% macro recall. This value was similar to GeneDisco-CoreSet (8.8%) and GeneDisco-TopUncertain (9.3%), but SAIBO-LABO did not rank first on any task.

### 4.2 True Feedback Does Not Reliably Improve Perturbation Discovery

Figure 3 compares final recall under true, random, and no feedback. Across the six tasks, no agent showed a consistent improvement from true feedback. Relative to no feedback, the macro-averaged differences were 0.7, +1.8, +1.9, and +5.5 percentage points for BioDiscoveryAgent, Biomni, Codex, and Codex+strategy, respectively. Relative to random feedback, the corresponding differences were +0.3, +3.4, 2.5, and +5.0 points. None of the eight 95% intervals excluded zero, and the contrast direction varied across agent–task pairs.

**Figure 3:**
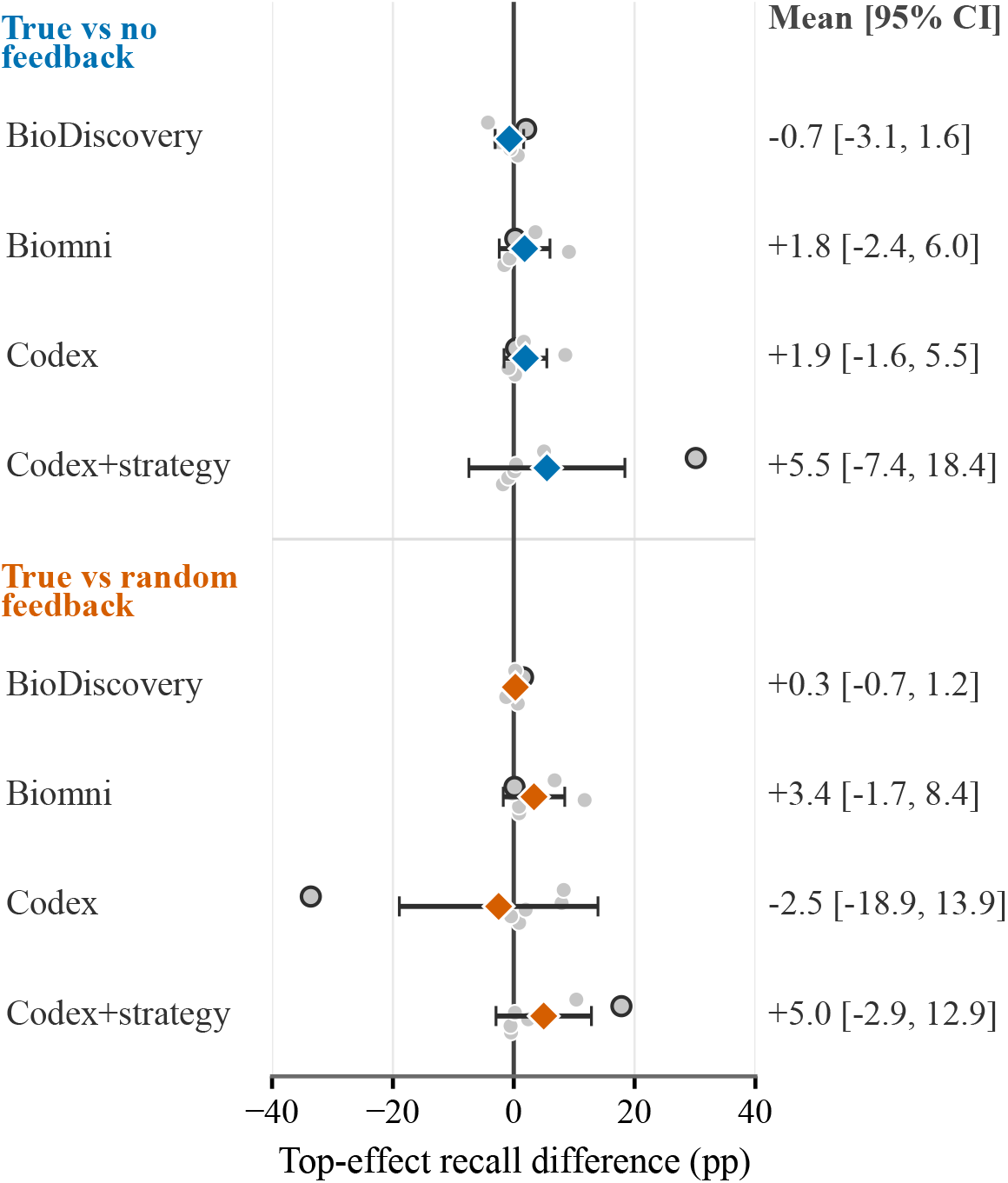
True-feedback contrasts in final recall. Gray circles show six task-level contrasts; outlined circles mark the phagocytosis task. Diamonds show task-macro means and whiskers show 95% *t* intervals across tasks. Positive values favor true feedback.

The apparent gain for Codex+strategy was largely driven by the phagocytosis task. On this task, true feedback exceeded no feedback by 30.1 points and random feedback by 17.8 points. After excluding this task, the two macro-averaged gains decreased from +5.5 to +0.6 points and from +5.0 to +2.4 points. For Codex, the true-versus-random difference changed from 2.5 to +3.7 points.

These results provided no evidence of a robust cross-task improvement from true feedback. Final recall, however, captures only the cumulative outcome after five rounds. It does not reveal how agents respond to individual feedback signals. We therefore examined whether feedback informed the next perturbation choices and whether those choices improved the subsequent outcome.

### 4.3 Feedback-Informed State Updates Rarely Guide Subsequent Actions

PerturbTrace tracks each transition along the Feedback–State–Action–Outcome sequence. It assesses whether the agent expresses a feedback-supported state update and whether the next batch follows that update. It also tests whether the batch finds more hits than expected under random sampling. Independent GPT-5.5 evaluations showed substantial agreement for Feedback-to-State and State-to-Action (Cohen *κ* = 0.715 and 0.723), supporting annotation reliability. Each feedback condition contains 288 transitions (four agents six tasks three seeds four transitions). Figure 4 summarizes the resulting stage-wise cascade under true and random feedback.

**Figure 4:**
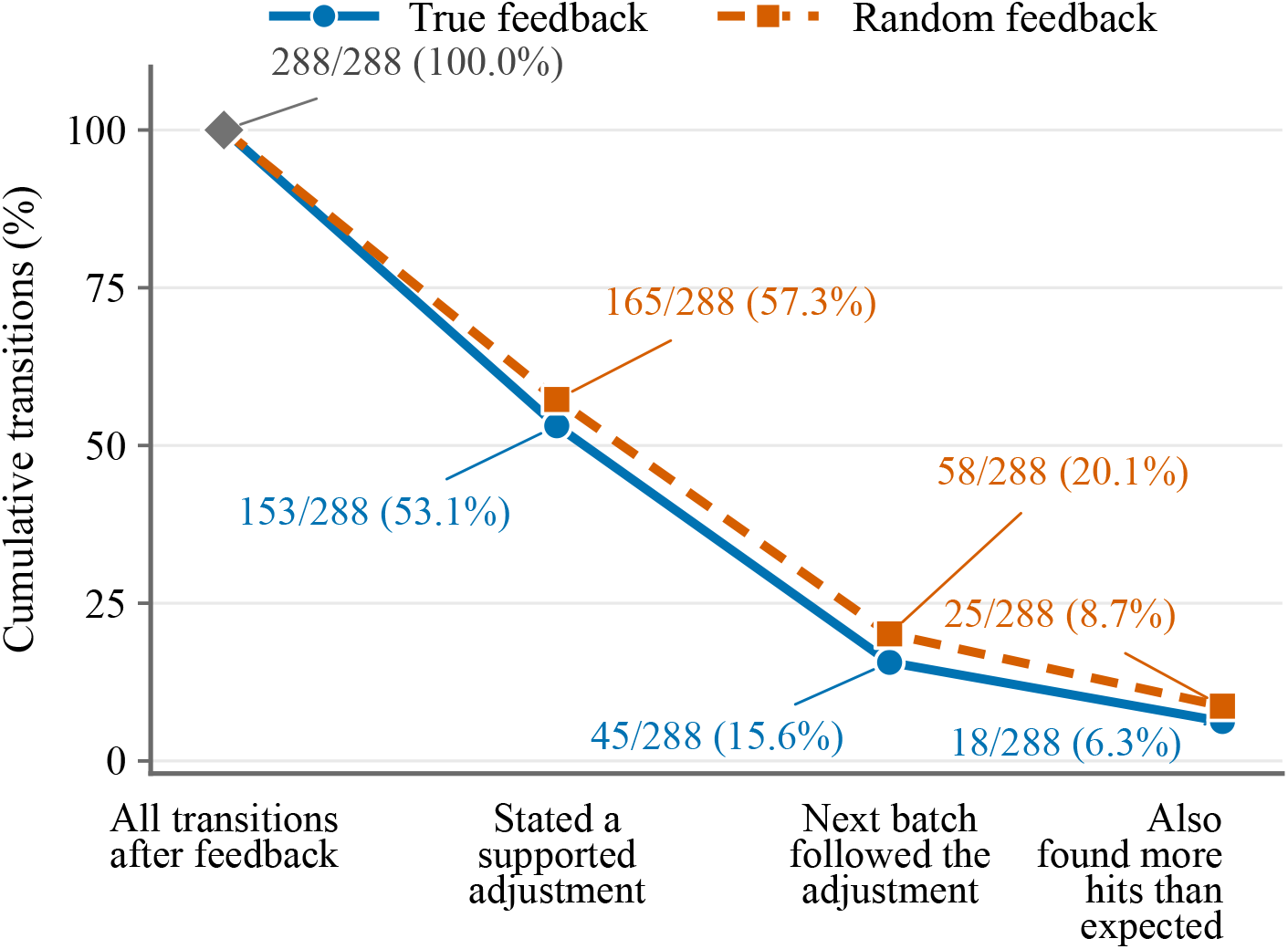
Feedback-to-outcome sequences. Each condition contributes 288 transitions. Later stages are subsets of earlier stages, and all percentages use 288 as the denominator. The final stage requires more benchmark-defined hits than expected under a hypergeometric baseline over the remaining pool. Random feedback reports counts after permuting the hidden action–outcome mapping.

Under true feedback, 153/288 transitions (53.1%) contained a feedback-supported state update.

Of all transitions, 45 (15.6%) carried this update into the next action, and 18 (6.3%) also achieved an above-random outcome. Under random feedback, the corresponding counts were 165 (57.3%), 58 (20.1%), and 25 (8.7%). Overall, only 43/576 transitions (7.5%) completed the full Feedback–State– Action–Outcome sequence. Because 25 occurred under random feedback, sequence completion alone did not establish the use of informative feedback. Table 1 reports the corresponding true-feedback cascade by agent.

**Table 1:** Cumulative true-feedback transitions by agent. Each agent contributes 72 transitions; later columns are subsets of earlier ones. Best values are bolded and second-best values are underlined.

| Agent | Supported | Implemented | Above exp. |
| --- | --- | --- | --- |
| BioDiscoveryAgent | 29 (40.3%) | 8 (11.1%) | 2 (2.8%) |
| Biomni | 24 (33.3%) | 12 (16.7%) | **7 (9.7%)** |
| Codex | 44 (61.1%) | 11 (15.3%) | 3 (4.2%) |
| Codex+strategy | **56 (77.8%)** | **14 (19.4%)** | <u>6 (8.3%)</u> |

We next examined where the sequence stopped under true feedback. Among 135 transitions without a supported state update, 80 contained interpretations unsupported by the aggregate feedback. Another 20 contained no identifiable feedback-related update, and 35 provided insufficient evidence for assessment. Among the 153 supported updates, 33 were partially reflected in the next batch, 32 conflicted with the accepted actions, and 43 were indeterminate. Only 45 updates were fully reflected in the accepted batch. Among these 45 transitions, 18 yielded above-random outcomes, whereas 27 did not. Thus, even supported and implemented updates did not necessarily improve immediate discovery.

Transition patterns varied across agents and tasks. For every agent, feedback-supported state updates occurred more often than implementation in subsequent actions. This pattern identified a consistent gap between state revision and action selection. Task-level cascades also differed substantially. The NK-cell exposure task progressed from 48 transitions to 24 supported updates, 1 implemented action, and 0 above-random outcomes. Project DRIVE retained 31 supported updates, 8 implemented actions, and 7 above-random outcomes.

### 4.4 Different Paths Lead to Similar Discovery Performance

Transition analysis showed that final recall alone could not determine whether feedback informed subsequent decisions. We therefore examined representative Project DRIVE runs that achieved similar final performance through different feedback-to-outcome paths.

One true-feedback Biomni run reached 46.6% recall through a complete Feedback–State–Action– Outcome sequence. After the hit count decreased from 65 to 28, the agent shifted the search from depleted core dependencies toward remaining viability genes. The subsequent batch reflected this adjustment and identified 43 hits, compared with 5.07 expected under random sampling.

In contrast, a true-feedback BioDiscoveryAgent run achieved higher recall (51.1%) without completing the sequence. Its rationales attributed the aggregate counts to RNA-processing and core-fitness mechanisms that the counts did not directly support. This trajectory showed that strong discovery performance can occur without explicit evidence that feedback guided the accepted actions.

Interestingly, a BioDiscoveryAgent run under random feedback also achieved high recall (48.1%). The displayed counts (5, 5, 8, and 2) were disconnected from the actual batch outcomes (71, 33, 31, 34, and 23 hits). Nevertheless, the agent produced coherent responses and selected effective perturbation batches. This case indicated that plausible feedback responses did not necessarily imply that the agent had extracted useful information from that feedback.

Finally, a low-performing BioDiscoveryAgent run illustrated the opposite scenario. After identifying 5 hits among 128 perturbations, the agent expanded the search in the next round without attributing the counts to a specific mechanism. Although the accepted batch implemented this adjustment, it yielded only 3 hits, compared with 5.79 expected under random sampling. The run ended with 17 hits (5.3% recall).

Together, these cases separated discovery performance, observable responses to feedback, and immediate experimental gains. High recall did not necessarily coincide with a complete observable sequence, and a feedback-consistent action did not guarantee an improved outcome.

## 5 Discussion and Limitations

One consistent reading of our results is that endpoint discovery and feedback use are different abilities, and the first does not require the second on these tasks. The seventeen tasks are screen-derived, and their hidden hit sets are effect thresholds on public measurements. An agent can score well by recovering perturbation-effect associations that already appear in the literature, in pretraining corpora, or in public resources, without changing its next-round behavior in response to the feedback it receives. This bears on what the benchmark measures. A genuinely closed-loop discovery agent faces a harder test: it should learn from experimental data that is out of distribution relative to its training, and it should turn that data into hypotheses on a new task where prior knowledge is silent. Our current tasks do not force that situation, because the answer is often recoverable from priors. The low full-chain rate is therefore not a general verdict on these agents. It is a measurement of feedback use under tasks that may not demand it, and it points to a concrete next step: build tasks whose hit sets cannot be recovered from public priors, so that feedback becomes necessary rather than optional.

A second question is what feedback use should mean for an in-context agent. In active learning and Bayesian optimization the term is precise: observed outcomes update a model or an acquisition function. An LLM agent performs no such explicit update, so its feedback use can appear only in what it writes and what it selects next. We operationalize feedback use as a three-stage chain, from a feedback-supported rationale, to a batch that implements that rationale, to an outcome better than random. This is a concrete and auditable definition, and it is a necessary step toward testing a behavior the field has mostly asserted. It is not the only definition. An agent may use feedback to confirm its current direction and change nothing, which the calibrated-update label allows for but which observable behavior cannot fully separate from ignoring feedback. Credit assignment, the step that decides which tested action caused a hit, is internal and invisible to the trace. These limits mark the boundary of what the framework can claim; the contribution is a working definition that can be tested, extended, and replaced, not a finished theory of feedback use.

The trace labels are themselves produced by language models. Two independent AI judges code whether a rationale is supported by feedback and whether the accepted batch implements it, with Cohen’s *κ* around 0.72. Agreement between the judges shows the coding is reproducible; it does not by itself establish validity. We do not treat this as a benchmark failure. Human experts would add their own, less scalable biases, and the practical alternative to automated coding is to abandon process measurement altogether. The property should stay visible, because any downstream use of the trace, including using it to train better agents, inherits this measurement.

The most actionable finding is where the chain breaks. Agents often write rationales that the judge marks as feedback-supported, but they rarely carry those rationales into the accepted batch. The fine-grained experiment in the appendix makes the point sharper: returning per-gene scores for tested candidates raises the number of stated adjustments while implementation stays near zero, and final recall does not improve. The bottleneck is not feedback perception or feedback granularity. It is the mechanism that turns a stated rationale into a concrete set of actions. If the goal is to build agents that learn from experiments, the leverage is in that action-selection layer, for example in explicit strategies that force each accepted batch to follow a stated direction, rather than in supplying more or finer feedback.

## 6 Conclusion

PerturbTrace reveals that discovery performance and observable feedback use are distinct aspects of closed-loop perturbation discovery. Across 17 screen-derived tasks, LLM agents achieved strong discovery performance. However, true feedback did not consistently improve final recall over random or no feedback. Only 43 of 576 feedback-bearing transitions completed the observable Feedback–State– Action–Outcome sequence, including 25 under random feedback. These results show that endpoint success alone does not establish effective feedback-guided adaptation. PerturbTrace provides a reusable framework for evaluating both what scientific agents discover and whether experimental evidence is reflected in subsequent experimental choices.

## Data and Code Availability

The PerturbTrace framework, PerturbTraceBench task definitions, and evaluation code used in this study are publicly available at https://github.com/HSZD-Team/PerturbTrace.

The underlying perturbation-screen datasets are available from the original public sources described in Appendix A.3.

## A Benchmark Construction and Task Provenance

### A.1 Task Contract and Information Boundary

A task is 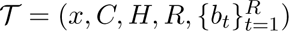, where *x* is the public scientific objective, *C* is the task-specific candidate space, *H ⊂ C* is the hidden hit set, *R* = 5, and *b_t_* is the round-specific batch size. The agent receives *x*, the budget, the previously submitted identifiers, and the identifiers in *C*. It does not receive hidden scores, hit labels, global ranks, source thresholds, or candidate-level outcomes. The evaluator retains the hidden action–outcome mapping and uses it for feedback generation and scoring.

For a source with multiple cell lines or assay contexts, one context is frozen per task. GDSC2, DepMap/Achilles, and Project DRIVE use the context with the greatest non-null candidate coverage in the frozen source release: HT-29, OVCAR-8, and SF-295, respectively. JUMP Cell Painting uses U2OS CRISPR profiles, and the Wang screen uses KBM7 cells (Iorio et al. 2016; Tsherniak et al. 2017; McDonald et al. 2017; Chandrasekaran et al. 2023; Wang et al. 2014).

### A.2 Frozen Tasks

Tables 2 and 3 give the source and frozen hit rule for each task. *N* is the candidate count and *H* is the hidden-hit count. Source identifiers refer to the release artifact. Benchmark membership is determined by the frozen hit rule.

**Table 2:** Frozen task provenance, T01–T09. BioDiscoveryAgent tasks are imported from the BioDiscoveryAgent task suite (Roohani et al. 2025).

| ID | Source, release, and context | Perturbation/readout | $N$ | $H$ | Frozen hit rule | Budget |
| --- | --- | --- | --- | --- | --- | --- |
| T01 | BioDiscoveryAgent S03, Carnevale22; engineered T cells under adenosine-like immunosuppression | KO; proliferation change | 18,861 | 943 | Directional efficacy boost or large absolute effect, frozen before formal runs | 5×128 |
| T02 | BioDiscoveryAgent S07, Steinhart CRISPRa; HA GD2 CAR, day 22 | CRISPRa; normalized sgRNA log-fold change | 18,797 | 149 | Top positive resistance/exhaustion-rescue score | 5×128 |
| T03 | ORCS 2.0.18, Screen 1953; Hagel, PMID 36548402; HUPT3 under CAR-T exposure | KO; exposure resistance | 20,079 | 884 | Source log2 fold change > 0.5 | 5×128 |
| T04 | ORCS 2.0.18, Screen 1954; Hagel, PMID 36548402; KP4 under CAR-T exposure | KO; exposure resistance | 20,079 | 532 | Source log2 fold change > 0.5 | 5×128 |
| T05 | JUMP cpg0016 assembled CRISPR profiles v1.0a; U2OS | CRISPR KO; 50-PC morphology shift | 7,972 | 399 | Top 5% by absolute morphology-shift score | 5×64 |
| T06 | BioDiscoveryAgent S06, Scharenberg22; choline-limitation assay | KO; sgRNA enrichment/depletion | 1,055 | 53 | High positive score or large absolute score after assay review | 5×32 |
| T07 | Wang et al., PMID 24336569; publication Dataset 1; KBM7 | KO; survival/enrichment | 7,114 | 321 | Corrected $p < 0.05$, represented by transformed score > 1.30103 | 5×128 |
| T08 | DepMap Public 26Q1; OVCAR-8 (ACH-000696) | KO; dependency/fitness | 18,513 | 926 | Top 5% dependency score | 5×128 |
| T09 | GDSC release 8.5, GDSC2 fitted response 27Oct23; HT-29 (SIDM00136) | Small molecule; sensitivity z-score | 286 | 15 | Top 5% sensitivity score | 5×32 |

**Table 3:** Frozen task provenance, T10–T17. The T12 rule is the formal benchmark rule; the source publication’s author-level *p*-value label is not used as the benchmark hit set.

| ID | Source, release, and context | Perturbation/readout | $N$ | $H$ | Frozen hit rule | Budget |
| --- | --- | --- | --- | --- | --- | --- |
| T10 | BioDiscoveryAgent S01, Schmidt-family primary T-cell screen; Science DOI 10.1126/science.abj4008 | Loss of function; normalized IFNG production | 18,418 | 920 | Top 5% by absolute score | 5×128 |
| T11 | BioDiscoveryAgent S02, same Schmidt-family screen | Loss of function; normalized IL2 production | 18,939 | 654 | Top 5% by absolute score | 5×128 |
| T12 | ORCS 2.0.18, Screen 1081; Zhuang et al., PMID 31921143; activated NK-cell exposure | KO; exposure sensitivity | 20,617 | 1,031 | Top 5% in the sensitive direction in the frozen formal task | 5×128 |
| T13 | ORCS 2.0.18, Screen 1140; Haney et al., PMID 30397336 | KO; magnetic-selection phagocytosis phenotype | 20,398 | 260 | CasTLE score > 21.2 | 5×128 |
| T14 | FITdb API snapshot, accessed 2026-06-10; Shifrut et al., PMID 30449619 | KO; primary T-cell proliferation phenotype | 19,106 | 962 | Frozen advantageous phenotype score < 0.05 | 5×128 |
| T15 | Project DRIVE, DEMETER2 Data v6 (2020-04-09); SF-295 | RNAi knockdown; dependency/viability | 7,975 | 399 | Top 5% dependency score | 5×128 |
| T16 | BioDiscoveryAgent S04, Sanchez21; neuronal model | KO; endogenous tau abundance | 18,469 | 924 | Top 5% by absolute tau change | 5×128 |
| T17 | BioDiscoveryAgent S05, directional variant of S04; neuronal model | KO; tau-lowering abundance change | 18,469 | 924 | Frozen directional tau-lowering compatibility set | 5×128 |

### A.3 Source Access and Preprocessing

**Table 4:** Recorded source access and preprocessing for non-BioDiscoveryAgent tasks.

| Tasks | Frozen source | Access handle | Preprocessing entry point |
| --- | --- | --- | --- |
| T03, T04, T12, T13 | BioGRID ORCS 2.0.18 static human-screen archive | https://downloads.thebiogrid.org/Download/BioGRID-ORCS/Latest-Release/BIOGRID-ORCS-ALL-homo_sapiens-LATEST.screens.tar.gz | `prepare_orcs_static_screen_task_table.py`, followed by task normalization |
| T05 | JUMP cpg0016 CRISPR v1.0a profiles | https://cellpainting-gallery.s3.amazonaws.com/cpg0016-jump-assembled/source_all/workspace/profiles_assembled/CRISPR/v1.0a/profiles_wellpos_cc_var_mad_outlier_featselect_sphering_harmony_PCA_corrected.parquet | `prepare_jump_cell_painting_task_table.py` |
| T07 | Wang Science 2014 supplementary Table 4 | https://orcs.thebiogrid.org/uploads/processed/5942c003b8b3f/24336569%20S%20Table%204.xlsx | `prepare_wang_crispr_task_table.py` |
| T08 | DepMap Public 26Q1 | https://depmap.org/portal/api/download/files | `normalize_action_score_table.py` after context selection |
| T09 | GDSC2 fitted dose response, release 8.5 | https://cog.sanger.ac.uk/cancerrxgene/GDSC_release8.5/GDSC2_fitted_dose_response_27Oct23.xlsx | `normalize_action_score_table.py` after context selection |
| T14 | FITdb static API snapshot | https://fitdb.lji.org/api/cell_vs_cell_4/advantageous | `normalize_action_score_table.py` |
| T15 | DEMETER2 Data v6 | https://ndownloader.figshare.com/files/11489693 | Context extraction followed by `normalize_action_score_table.py` |

### A.4 Feedback-Condition Panel

The feedback-condition panel contains six gene-level tasks with five rounds, 128 actions per round, and a total budget of 640 actions. Candidate-space coverage is 640*/N*, and hit prevalence is *H/N* . The task-property strata were fixed before the formal runs.

**Table 5:** Prespecified six-task feedback-condition panel.

| Stratum | Task | Context | $N$ | $H$ | Coverage | Prevalence |
| --- | --- | --- | --- | --- | --- | --- |
| Budget-easy | T07 | KBM7 CRISPR selection | 7,114 | 321 | 9.00% | 4.51% |
| Budget-easy | T15 | SF-295 Project DRIVE RNAi dependency | 7,975 | 399 | 8.03% | 5.00% |
| Standard genome-scale | T03 | HUPT3 CAR-T exposure resistance | 20,079 | 884 | 3.19% | 4.40% |
| Standard genome-scale | T12 | NK-cell exposure CRISPR screen | 20,617 | 1,031 | 3.10% | 5.00% |
| Feedback-sparse | T02 | GD2 CAR-T exhaustion-resistance CRISPRa | 18,797 | 149 | 3.40% | 0.79% |
| Feedback-sparse | T13 | Phagocytosis CRISPR screen | 20,398 | 260 | 3.14% | 1.27% |

## B Execution and Feedback Protocol

**Figure 5:**
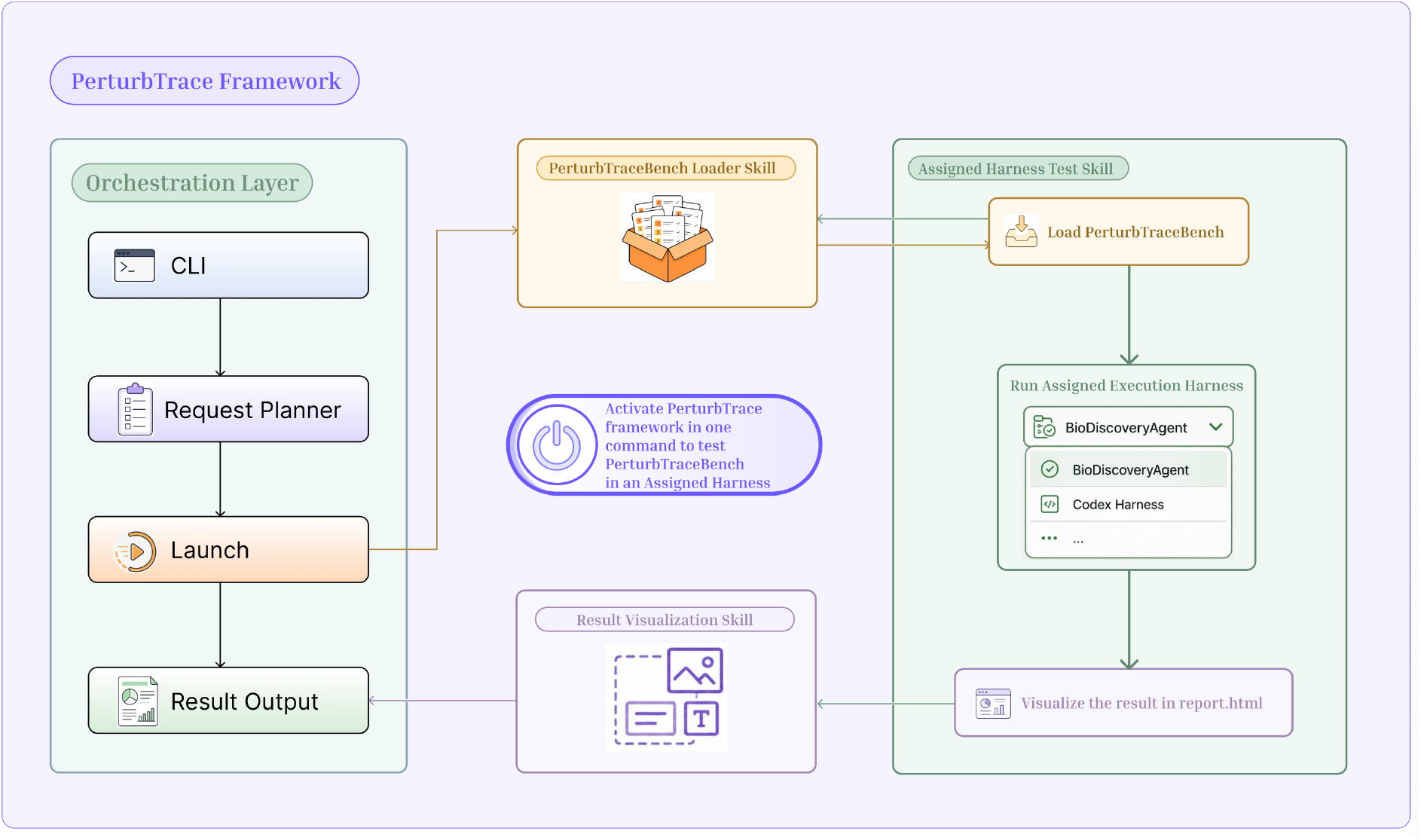
PerturbTrace execution workflow. The orchestration layer receives a command, constructs a request plan, launches the benchmark loader and assigned agent harness, validates the submitted batch, queries the hidden oracle, and writes machine-readable run artifacts. The result-visualization component renders reports from these artifacts.

### B.1 Round-Level Execution

Each run has its own task–agent–feedback-policy–seed configuration. At the start of a round, the harness loads the public task brief, the untested candidate identifiers, prior accepted actions, and any feedback allowed by the assigned condition. It invokes the agent through its frozen adapter and parses the scientific rationale and submitted `Solution:` field. The submitted batch is then checked for membership, uniqueness, prior use, and exact size. If the batch is invalid, the harness requests an action-only membership or format repair. Once validation succeeds, the harness freezes the accepted batch, queries the hidden oracle, updates the feedback state for the assigned condition, and writes the trace and round artifacts.

#### Repair semantics

A repair only makes the submitted action list executable. It replaces invalid or previously used identifiers, removes duplicates, and fills a short batch to the required size. The repair prompt asks for one revised `Solution:` line without scientific prose, so the original rationale and search direction remain unchanged. The State-to-Action audit compares that rationale with the final accepted batch from the same round. Repair does not alter the hidden action–outcome mapping or the delivered feedback.

### B.2 Feedback Conditions

**Table 6:**
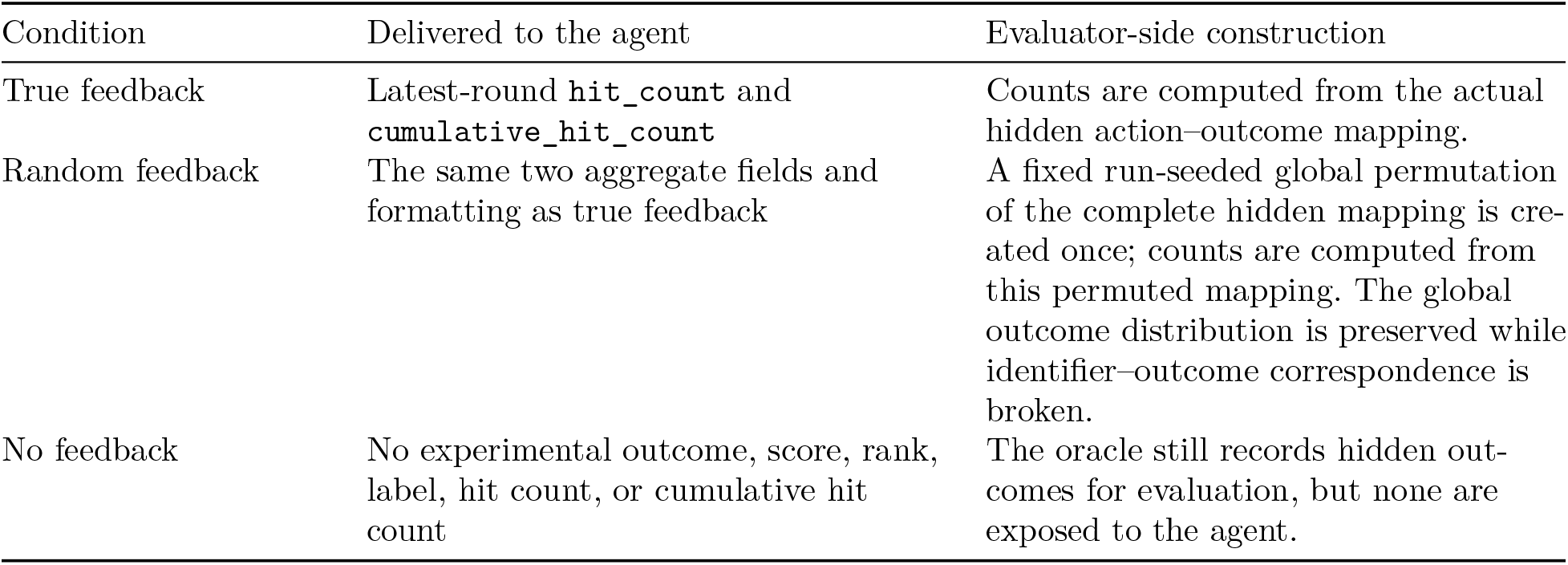
Information delivered under each feedback policy.

| Condition | Delivered to the agent | Evaluator-side construction |
| --- | --- | --- |
| True feedback | Latest-round `hit_count` and `cumulative_hit_count` | Counts are computed from the actual hidden action-outcome mapping. |
| Random feedback | The same two aggregate fields and formatting as true feedback | A fixed run-seeded global permutation of the complete hidden mapping is created once; counts are computed from this permuted mapping. The global outcome distribution is preserved while identifier-outcome correspondence is broken. |
| No feedback | No experimental outcome, score, rank, label, hit count, or cumulative hit count | The oracle still records hidden outcomes for evaluation, but none are exposed to the agent. |

Counts are computed from the actual hidden action–outcome mapping.

A fixed run-seeded global permutation of the complete hidden mapping is created once; counts are computed from this permuted mapping. The global outcome distribution is preserved while identifier–outcome correspondence is broken.

The oracle still records hidden outcomes for evaluation, but none are exposed to the agent.

Random feedback is fixed within a run. All Action-to-Outcome analyses use the actual hidden hit set, including for random-and no-feedback runs. Thus, displayed random counts affect the agent-facing interaction but never redefine evaluator-side productivity.

### B.3 IFNG Feedback-Granularity Audit Protocol

The IFNG audit compares two interfaces over five completed five-round trajectories per interface. Fine feedback returns per-gene/signal results for candidates tested in the preceding batch. True feedback returns the aggregate `hit_count` and `cumulative_hit_count` fields used in the main experiment. The same three-stage labels defined in Section D are reported for the resulting 20 adjacent-round transitions per interface.

### B.4 Agent Configurations

Four agent configurations are evaluated. Each configuration contributes 51 true-feedback runs over the 17 tasks and 18 runs under each additional condition over the six-task panel. All four configurations use `xhigh` reasoning effort.

**Table 7:** Frozen agent configurations.

| Configuration | Interface | Model / effort | Tool and candidate-delivery settings | Additional procedure |
| --- | --- | --- | --- | --- |
| BioDiscoveryAgent | Streaming Responses adapter with draft critique | GPT-5.5 / **xhigh** | Web search disabled; task-declared identifiers-only artifact delivered once through the no-tool transport | Biomedical discovery workflow |
| Biomni | Biomni A1 | GPT-5.5 / **xhigh** | Tool retriever enabled; no expected data-lake files; complete identifiers-only list in the initial prompt and isolated artifact |  |
| Codex | Codex CLI | GPT-5.5 / **xhigh** | Web search enabled | Native-neutral, task-neutral execution skill |
| Codex+strategy | Codex CLI | GPT-5.5 / **xhigh** | Web search enabled | Restricted perturbation-selection strategy |

### B.5 Sequential Baselines and Frozen Hyperparameters

Random samples uniformly without replacement from the remaining candidate pool using the run seed. GeneDisco CoreSet and TopUncertain use the GeneDisco 1.0.5 implementation (Mehrjou et al. 2022). Both receive native per-action signed outcomes, retrain a single-hidden-layer MC-dropout MLP from scratch after each round, and use STRING Mashup features. The first feature column is dropped for package parity, leaving 799 columns, followed by a fixed 64-dimensional Gaussian projection with seed 2022. Unmapped identifiers use a deterministic SHA-256-counter Rademacher fallback.

SAIBO-LABO uses SAIBO 1.0.0, the same 64-dimensional STRING descriptor, and GPT-5.5 at xhigh effort for online low-fidelity predictions. Its response limit is 12,000 tokens, timeout is 480 seconds, and the transport permits seven retries with 10–60 second backoff. Temperature and top_p were not set. The SAIBO configuration was frozen before its canary run.

### B.6 Run Accounting and Randomness

Seeds 1, 2, and 3 control harness sampling, random-baseline sampling, and the run-specific global permutation used for random feedback.

**Table 8:** Frozen baseline settings.

| Method | Acquisition and model | Final settings |
| --- | --- | --- |
| Random | Uniform sampling without replacement | Seeds 1, 2, and 3; task-specific batch size; five rounds |
| GeneDisco CoreSet | Greedy max–min distance in the learned hidden representation | Hidden size 32; dropout 0.1; learning rate 0.01; maximum 100 epochs; validation fraction 0.2; early-stopping patience 10; 64 projected features |
| GeneDisco TopUncertain | Highest predictive standard deviation among untested actions | Hidden size 32; dropout 0.5; 100 consistent MC-dropout samples; learning rate 0.01; maximum 100 epochs; validation fraction 0.2; patience 10; 64 projected features |
| SAIBO-LABO | Finite-pool LABO adapter; low-fidelity LLM predictions fused as $\rho$ times a low-fidelity GP plus a discrepancy GP; UCB acquisition | UCB $\beta = 1.0$; mismatch threshold 0.75; force high-fidelity after two low-fidelity steps; at most two low-fidelity loops; shortlist multiplier 2, min 64, max 256; low-fidelity exploration multiplier 2; at most 1,024 low-fidelity training points; five GP iterations; LLM batch 128; two parse retries |

**Table 9:**
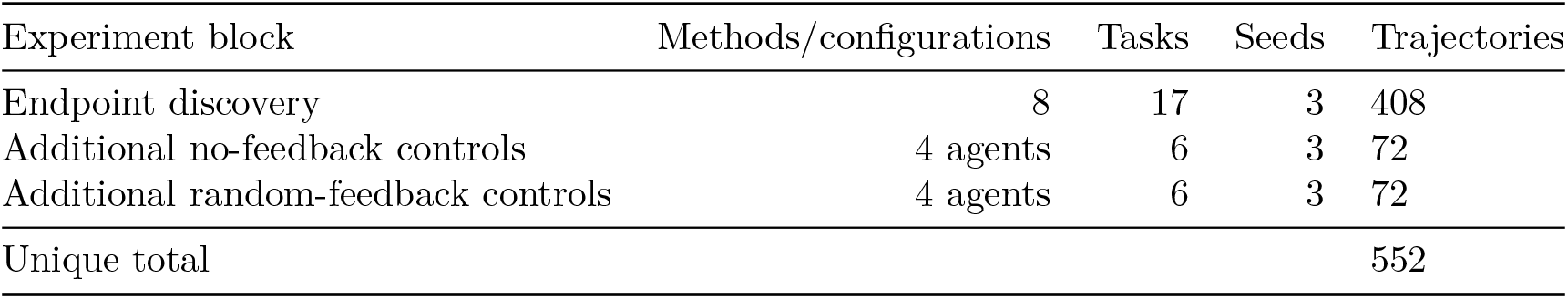
Unique formal trajectories used in the paper. The 72 true-feedback panel trajectories are already included in each agent’s 17-task endpoint runs and are not counted twice.

## C Metrics and Statistical Analysis

### C.1 Endpoint Discovery

For task *i*, let *H_i_* be its hidden hit set and 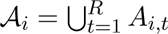 be all accepted actions. Final top-effect recall is

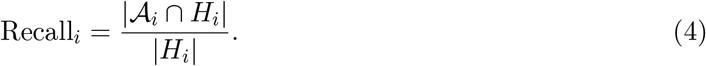

This metric is used because the experimental objective is to recover benchmark-defined high-effect perturbations under a fixed budget. The endpoint benchmark reports the mean and sample standard deviation over three seeds for each task–method cell. Macro recall is the unweighted mean of the 17 task means.

### C.2 Feedback-Condition Contrasts

Within every agent–task–condition cell, recall is first averaged over the three seeds. For each agent, the six task-level differences 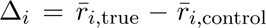 are then macro-averaged. A two-sided 95% Student-*t* interval is

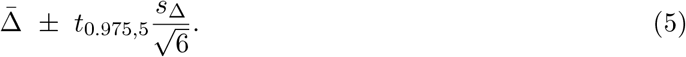

Tasks are the units for these intervals.

### C.3 Immediate Outcome

Before round *t* + 1, let *N_t_* and *H_t_* be the remaining candidate and hidden-hit counts after removing all prior accepted actions. For accepted batch size *b_t_*_+1_ and true hit count *h_t_*_+1_, RHE is

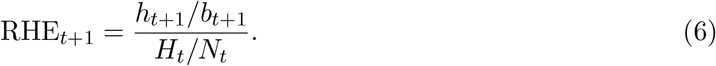

With *X_t+1_~* Hypergeom(*N_t_, H_t_, b_t_*_+1_), an outcome is classified as above random expectation when *P* (*X_t_*_+1_ ≥ *h_t_*_+1_) ≤ 0.025, below random expectation when *P* (*X_t_*_+1_ ≤ *h_t_*_+1_) ≤ 0.025, and within random expectation otherwise. If *H_t_* = 0, RHE is undefined. All policies are scored against the actual hidden hit set.

### C.4 Process Rates

Wilson score intervals are reported for transition proportions. The calibrated-update denominator is all feedback-bearing transitions. The diagnostic action-translation denominator is all assessable transitions with an established state update (calibrated or over-attributed), with `aligned` as the numerator. Outcome evaluation uses all oracle-computable transitions. Full-chain completion uses all feedback-bearing, oracle-computable transitions and requires a calibrated update, aligned action, and outcome above random expectation. For the paper’s cascade, the State-to-Action denominator is restricted to calibrated updates. The later columns are therefore literal subsets of the earlier columns.

## D Transition Audit Protocol

### D.1 Audit Unit and Observable State

Each five-round trajectory contributes four transitions,

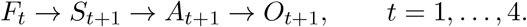

*S_t_*_+1_ contains only the scientific content recorded in the submitted rationale, including hypotheses or mechanisms, the agent’s stance toward the evidence, stated uncertainty, and its next-round search strategy. We do not infer missing content from the accepted action or outcome. An explicit reference such as “continue the previous core strategy” may be interpreted using the previous rationale, but an unstated carry-forward is not coded.

### D.2 Feedback-to-State

For aggregate feedback, a mechanistic claim is coded as over-attributed when the displayed counts do not support that claim.

### D.3 State-to-Action

State and action are coded independently. State coders see the public task context, previous rationale, delivered feedback, current rationale, and frozen task vocabulary. They do not see the current action, agent identity, condition label, or outcomes. Action composition is constructed from public MyGene.info names, summaries, GO biological-process terms, and a frozen task-specific primary-module map. It excludes rationale, feedback, agent identity, condition, and all oracle information.

**Table 10:** Feedback-to-State fields and composite rule. Rules are applied in the listed order. Random feedback is judged against the feedback delivered to the agent, never against hidden truth.

| Field | Values | Operational meaning |
| --- | --- | --- |
| Grounding | 0, 1, 2 | No link; ambiguous/partial link; explicit and substantive link between delivered feedback and the next decision |
| Operation | maintain/reinforce, expand, narrow/prune, pivot, mixed, none, unclear | Search-direction operation explicitly stated in the rationale |
| Calibration | calibrated, over-attributed, unclear | Whether the inference is supported by the information content of aggregate feedback |
| No feedback | `not_applicable_control` | Feedback-to-State is not defined |
| Grounding 0 or operation none | `no_observable_update` | No observable feedback-linked update |
| Grounding 1, unclear operation, or unclear calibration | `indeterminate` | Evidence is insufficient for a determinate label |
| Grounding 2, explicit operation, over-attribution | `over_attributed_update` | The rationale links feedback to a mechanistic claim not identified by aggregate counts |
| Grounding 2, explicit operation, calibrated inference | `calibrated_update` | Observable feedback-supported state update |

For a batch of 128, a module is absent at 0 genes, minor at 1–12, and major at 13 or more. A batch is unassessable when at least 39 actions map to `other_uncertain`. Maintain/reinforce requires retained modules not to decrease. Expand requires retained directions not to decrease and at least one priority-up module to increase. Narrow/prune requires at least one priority-down module to decrease and retained module concentration to increase. Pivot requires a priority-up module to become major and a previously major priority-down module to become minor or absent. Mixed operations must satisfy every explicitly stated component. Labels are `aligned, partially_aligned, mismatch`, and `unassessable`.

### D.4 Full-Chain Criterion

A transition is full-chain positive when and only when

`calibrated_update ∧ aligned ∧` outcome above random expectation.

This conjunction defines the full-chain criterion.

### D.5 Blinding, Double Coding, and Reliability

The audit was frozen on 2026-07-20 and reads completed run artifacts without modifying run roots. The 288 double-coded transitions contain one deterministic seed from every agent task condition transition-position stratum. Exactly 144 strata form the development pilot and 144 form held-out confirmation. Decision rules could be revised on development only. Coder A and coder B were independent GPT-5.5 medium-effort API passes with separate calls and reversed batch order. The prespecified gate required held-out Cohen’s *κ* ≥ 0.70 for the composite Feedback-to-State and State-to-Action labels.

**Table 11:** Frozen task-specific module vocabularies used in the six-task audit. Every vocabulary also contains other_uncertain. Full keywords and symbol patterns are stored in module_codebooks_v1.json.

| Task | Primary modules |
| --- | --- |
| T02 | T-cell state/transcription; cytokine/costimulation; antigen-receptor signaling; survival/cell death; proliferation/cell cycle; metabolism/mitochondria; stress/redox/proteostasis; autophagy/lysosome; trafficking/adhesion/cytoskeleton; chromatin/gene regulation |
| T03 | Antigen presentation/interferon; death execution/survival; immune synapse/surface ligands; cytokine/innate inflammation; growth signaling; adhesion/trafficking/cytoskeleton; chromatin/lineage transcription; stress/autophagy/proteostasis; metabolism/mitochondria; cell cycle/DNA repair |
| T07 | Cell cycle/mitosis; DNA replication/repair; transcription/chromatin; RNA splicing/translation; proteostasis/ubiquitin; metabolism/mitochondria; membrane trafficking/cytoskeleton; signal transduction; cell death/stress; immune/inflammation |
| T12 | Antigen presentation/interferon; NK ligands/immune synapse; death execution/survival; checkpoint/immunosuppression; cytokine/inflammation signaling; glycocalyx/adhesion/trafficking; stress/autophagy/lysosome; chromatin/lineage transcription; metabolism/mitochondria; cell cycle/DNA repair |
| T13 | Actin/cytoskeleton/motility; phagosome/endosome/lysosome; phagocytic receptors/opsonization; membrane trafficking/vesicle; adhesion/integrin/matrix; signaling/small GTPase; lipid metabolism/membrane; innate inflammation; transcription/chromatin; cell cycle/survival/stress |
| T15 | Cell cycle/mitosis; DNA replication/repair; transcription/chromatin; RNA splicing/translation; proteasome/ubiquitin/proteostasis; metabolism/mitochondria; oncogenic signaling; apoptosis/survival; membrane trafficking/cytoskeleton; immune/inflammation; lineage/development |

**Table 12:** Held-out agreement between coder A and coder B.

| Field | $n$ | Exact agreement | $\kappa$ | Weights |
| --- | --- | --- | --- | --- |
| Grounding | 96 | 89.6% | 0.859 | Quadratic |
| Operation | 144 | 81.9% | 0.758 | Unweighted |
| Calibration | 96 | 92.7% | 0.854 | Unweighted |
| Composite Feedback-to-State | 96 | 83.3% | 0.715 | Unweighted |
| State-to-Action | 144 | 79.9% | 0.723 | Unweighted |

## E Extended Results and Robustness Analyses

### E.1 Task-Level Endpoint Discovery

**Table 13:** Final top-effect recall on all 17 tasks. Entries are percentage means ± sample SD over three seeds. Uncertain denotes GeneDisco TopUncertain, and C+strat. denotes Codex+strategy.

| Task | Random | CoreSet | Uncertain | SAIBO | BioDiscoveryAgent | Biomni | Codex | C+strat. |
| --- | --- | --- | --- | --- | --- | --- | --- | --- |
| T01 | 3.8 ± 0.3 | 3.5 ± 0.5 | 3.7 ± 0.3 | 3.0 ± 0.5 | 4.3 ± 0.3 | 4.2 ± 0.2 | 7.3 ± 3.0 | 10.1 ± 5.1 |
| T02 | 4.9 ± 1.0 | 2.9 ± 0.4 | 3.1 ± 0.4 | 4.5 ± 0.4 | 13.0 ± 0.4 | 11.2 ± 0.8 | 11.2 ± 1.0 | 10.7 ± 1.2 |
| T03 | 3.5 ± 0.5 | 3.6 ± 0.4 | 3.7 ± 1.4 | 3.3 ± 0.8 | 5.9 ± 0.5 | 6.3 ± 0.3 | 6.0 ± 1.0 | 6.3 ± 0.4 |
| T04 | 3.5 ± 1.2 | 4.1 ± 0.8 | 4.3 ± 1.2 | 3.8 ± 1.7 | 5.5 ± 1.5 | 4.5 ± 0.5 | 4.9 ± 0.2 | 5.4 ± 0.1 |
| T05 | 4.1 ± 1.3 | 3.8 ± 0.6 | 7.3 ± 3.5 | 3.3 ± 0.3 | 11.9 ± 2.9 | 10.3 ± 0.7 | 16.2 ± 3.5 | 15.0 ± 0.7 |
| T06 | 15.7 ± 4.7 | 23.3 ± 2.2 | 16.4 ± 5.8 | 15.1 ± 3.3 | 39.6 ± 5.7 | 33.3 ± 4.7 | 81.8 ± 30.0 | 82.4 ± 30.5 |
| T07 | 7.2 ± 1.1 | 10.1 ± 1.5 | 15.3 ± 5.1 | 9.3 ± 1.9 | 7.4 ± 3.3 | 35.5 ± 3.3 | 10.8 ± 2.5 | 10.0 ± 1.1 |
| T08 | 3.7 ± 0.9 | 5.5 ± 1.7 | 12.3 ± 5.0 | 6.2 ± 0.1 | 47.6 ± 1.9 | 38.8 ± 0.8 | 45.4 ± 2.8 | 44.6 ± 2.2 |
| T09 | 62.2 ± 7.7 | 60.0 ± 6.7 | 51.1 ± 13.9 | 64.4 ± 10.2 | 64.4 ± 3.8 | 68.9 ± 3.8 | 68.9 ± 3.8 | 64.4 ± 3.8 |
| T10 | 3.5 ± 0.4 | 3.6 ± 0.2 | 4.2 ± 0.9 | 2.9 ± 0.5 | 10.8 ± 0.1 | 9.7 ± 0.2 | 20.8 ± 9.4 | 18.0 ± 14.4 |
| T11 | 2.7 ± 0.5 | 3.8 ± 0.5 | 6.5 ± 5.0 | 3.6 ± 0.6 | 11.5 ± 0.3 | 10.4 ± 0.5 | 10.9 ± 1.1 | 11.0 ± 0.6 |
| T12 | 2.8 ± 0.5 | 3.5 ± 0.6 | 3.5 ± 0.5 | 3.5 ± 0.5 | 4.1 ± 0.1 | 4.8 ± 0.4 | 12.9 ± 14.4 | 4.7 ± 0.8 |
| T13 | 2.1 ± 0.6 | 2.9 ± 0.2 | 3.6 ± 0.4 | 2.4 ± 0.6 | 14.5 ± 2.8 | 11.5 ± 1.3 | 11.4 ± 0.9 | 60.6 ± 2.6 |
| T14 | 3.7 ± 0.2 | 3.3 ± 0.4 | 6.1 ± 3.5 | 3.5 ± 0.7 | 7.8 ± 0.6 | 6.6 ± 0.5 | 6.9 ± 0.3 | 7.1 ± 0.3 |
| T15 | 7.9 ± 0.9 | 8.9 ± 1.3 | 9.9 ± 1.7 | 11.0 ± 0.7 | 47.2 ± 6.8 | 44.5 ± 1.8 | 44.4 ± 0.7 | 45.1 ± 1.1 |
| T16 | 3.3 ± 0.2 | 3.4 ± 0.3 | 4.0 ± 0.8 | 4.0 ± 0.9 | 4.4 ± 0.6 | 3.5 ± 0.5 | 28.4 ± 6.9 | 28.5 ± 2.1 |
| T17 | 3.0 ± 0.9 | 3.9 ± 1.5 | 3.7 ± 1.6 | 3.5 ± 0.3 | 7.2 ± 0.5 | 6.9 ± 0.5 | 26.0 ± 16.6 | 39.9 ± 1.8 |
| Macro mean | 8.1 | 8.8 | 9.3 | 8.7 | 18.1 | 18.3 | 24.4 | 27.3 |

### E.2 Task-Level Feedback Conditions

**Table 14:**
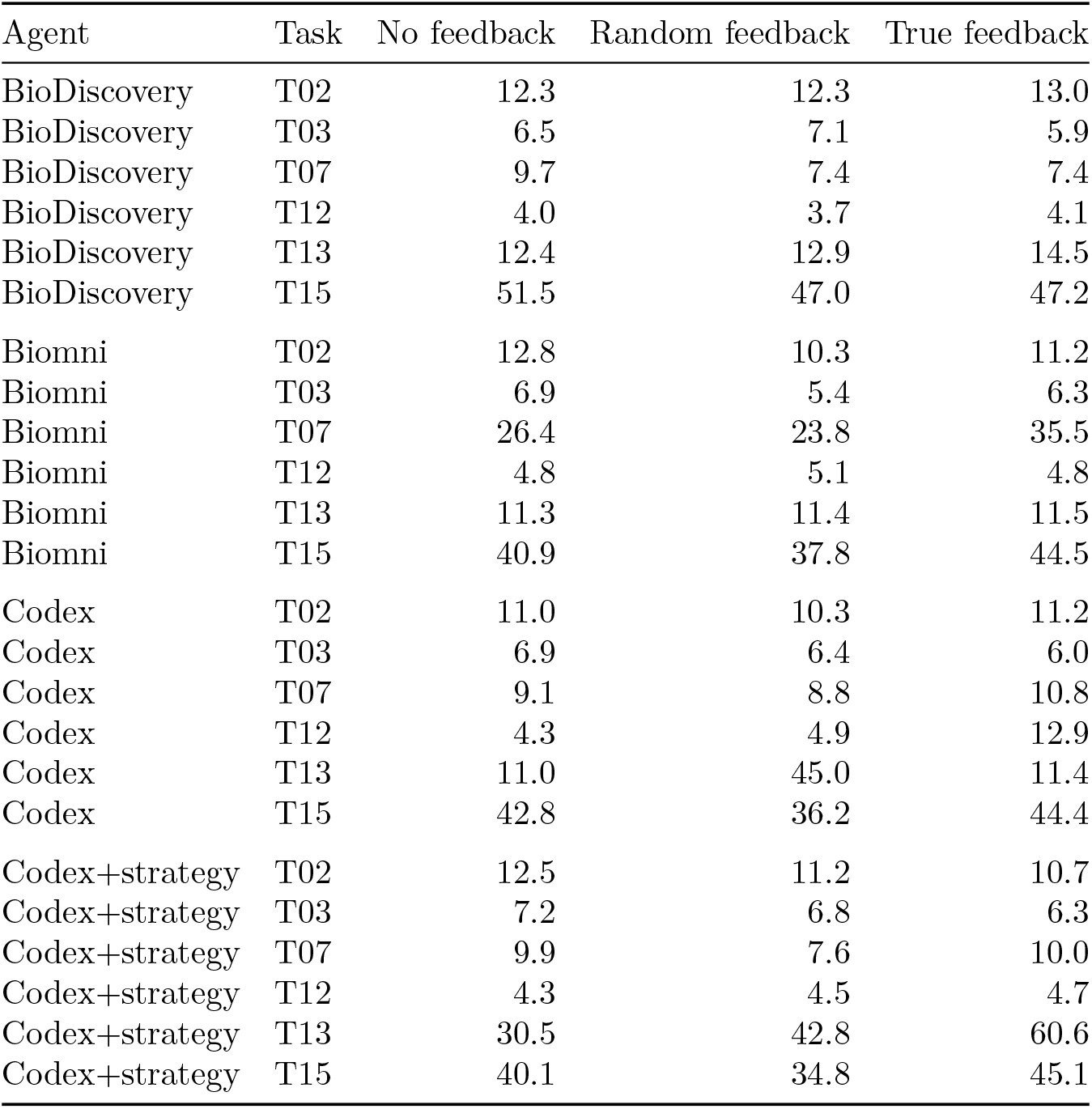
Final recall on every agent–task–feedback cell. Values are percentage means over three seeds.

**Table 15:**
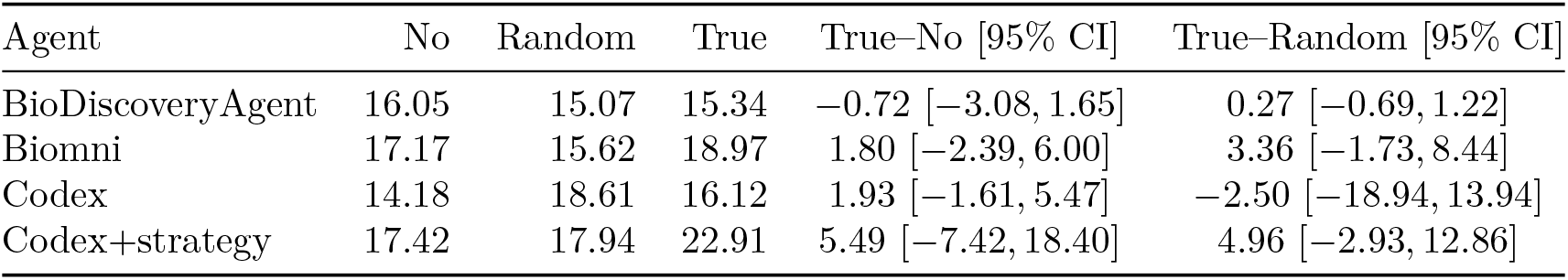
Six-task macro recall and true-feedback contrasts. Recall columns are percentages. Contrasts and 95% *t* intervals are percentage points across six task-level differences.

| Agent | No | Random | True | True–No [95% CI] | True–Random [95% CI] |
| --- | --- | --- | --- | --- | --- |
| BioDiscoveryAgent | 16.05 | 15.07 | 15.34 | −0.72 [−3.08, 1.65] | 0.27 [−0.69, 1.22] |
| Biomni | 17.17 | 15.62 | 18.97 | 1.80 [−2.39, 6.00] | 3.36 [−1.73, 8.44] |
| Codex | 14.18 | 18.61 | 16.12 | 1.93 [−1.61, 5.47] | −2.50 [−18.94, 13.94] |
| Codex+strategy | 17.42 | 17.94 | 22.91 | 5.49 [−7.42, 18.40] | 4.96 [−2.93, 12.86] |

### E.3 Transition-Level Results

**Table 16:** Condition-level process rates. Brackets are Wilson 95% intervals. Action translation is the diagnostic rate among assessable established updates; the cascade-aligned and full-chain columns use all 288 transitions as the displayed denominator.

| Condition | Calibrated update | Action translation | Above random expectation | Cascade aligned | Full chain |
| --- | --- | --- | --- | --- | --- |
| No feedback | NA | NA | 110/288, 38.2% [32.8,43.9] | NA | NA |
| Random | 165/288, 57.3% [51.5,62.9] | 85/188, 45.2% [38.3,52.4] | 102/288, 35.4% [30.1,41.1] | 58/288, 20.1% | 25/288, 8.7% [5.9,12.5] |
| True | 153/288, 53.1% [47.4,58.8] | 66/171, 38.6% [31.6,46.1] | 119/288, 41.3% [35.8,47.1] | 45/288, 15.6% | 18/288, 6.3% [4.0,9.7] |

**Table 17:** Complete transition flow under true and random feedback. State-to-Action rows are restricted to calibrated updates, so each block is a literal cascade.

| Stage | Label | True | Random |
| --- | --- | --- | --- |
| Feedback-to-State ($n = 288$) | Calibrated update | 153 | 165 |
|  | Over-attributed update | 80 | 69 |
|  | No observable update | 20 | 36 |
|  | Indeterminate | 35 | 18 |
| State-to-Action, given calibrated | Aligned | 45 | 58 |
|  | Partially aligned | 33 | 31 |
|  | Mismatch | 32 | 39 |
|  | Unassessable | 43 | 37 |
| Outcome, given calibrated and aligned | Above random expectation | 18 | 25 |
|  | Within random expectation | 27 | 33 |

**Table 18:** Complete cascades by agent. Every agent–condition row contains 72 transitions; later columns are subsets of earlier columns.

| Condition | Agent | Calibrated | Aligned | Above-random full chain |
| --- | --- | --- | --- | --- |
| True | BioDiscoveryAgent | 29 | 8 | 2 |
| True | Biomni | 24 | 12 | 7 |
| True | Codex | 44 | 11 | 3 |
| True | Codex+strategy | 56 | 14 | 6 |
| Random | BioDiscoveryAgent | 34 | 15 | 8 |
| Random | Biomni | 26 | 9 | 4 |
| Random | Codex | 47 | 16 | 7 |
| Random | Codex+strategy | 58 | 18 | 6 |

**Table 19:**
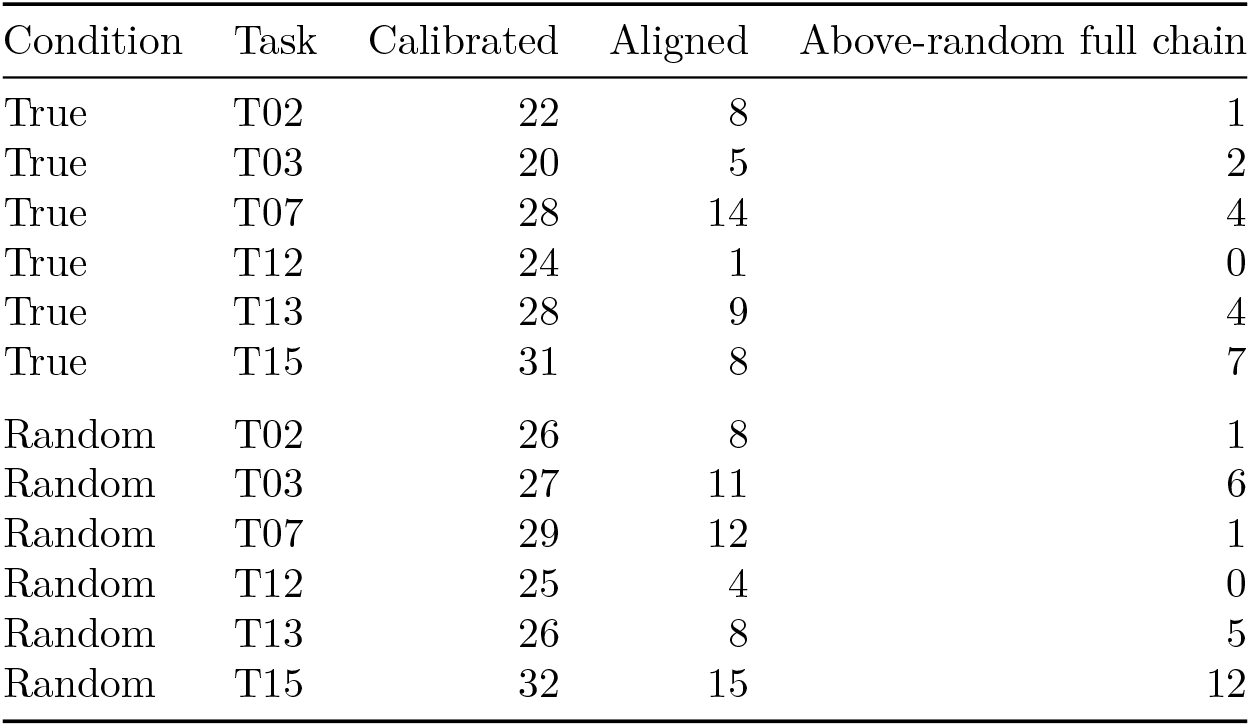
Complete cascades by task. Every task–condition row contains 48 transitions.

Across true and random feedback, 43/576 transitions completed all three stages: 18 under true feedback and 25 under random feedback.

### E.4 Action-to-Outcome Threshold Sensitivity

We evaluated the sensitivity of the full-chain rate to the upper-tail cutoff used in the Action-to-Outcome criterion. The analysis used the same 576 feedback-bearing transitions as the primary audit. For each transition, the one-sided hypergeometric test remained conditioned on the observed remaining candidate pool, remaining hidden-hit count, and accepted batch size. We changed only the cutoff for classifying an outcome as above random expectation, from *p* ≤ 0.025 to *p* ≤ 0.05 and *p* ≤ 0.10; all Feedback-to-State and State-to-Action labels were retained.

Table 20 reports the resulting counts. Under true feedback, full-chain completion increased from 18*/*288 (6.3%) at *p* ≤ 0.025 to 20*/*288 (6.9%) and 25*/*288 (8.7%) at the two relaxed cutoffs. Under random feedback, the corresponding counts were 25*/*288 (8.7%), 27*/*288 (9.4%), and 32*/*288 (11.1%). Across both conditions, full-chain completion was 43*/*576 (7.5%), 47*/*576 (8.2%), and 57*/*576 (9.9%), respectively. The full-chain rate under true feedback was not higher than that under random feedback at any cutoff.

**Table 20:** Sensitivity of the Action-to-Outcome threshold. Each transition uses a one-sided hypergeometric upper-tail probability conditioned on its actual remaining candidate pool, remaining hidden-hit count, and accepted batch size. Supported and Implemented are unchanged across thresholds. Full chain is reported over all 288 transitions per interface; Above random among implemented is reported over implemented transitions. Brackets give Wilson 95% intervals.

| Feedback interface | Upper-tail threshold | Supported | Implemented | Full chain | Above random among implemented |
| --- | --- | --- | --- | --- | --- |
| True feedback | 0.025 | 153/288 (53.1%) | 45/288 (15.6%) | 18/288 (6.3%; [4.0, 9.7]) | 18/45 (40.0%; [27.0, 54.5]) |
| True feedback | 0.050 | 153/288 (53.1%) | 45/288 (15.6%) | 20/288 (6.9%; [4.5, 10.5]) | 20/45 (44.4%; [30.9, 58.8]) |
| True feedback | 0.100 | 153/288 (53.1%) | 45/288 (15.6%) | 25/288 (8.7%; [5.9, 12.5]) | 25/45 (55.6%; [41.2, 69.1]) |
| Random feedback | 0.025 | 165/288 (57.3%) | 58/288 (20.1%) | 25/288 (8.7%; [5.9, 12.5]) | 25/58 (43.1%; [31.2, 55.9]) |
| Random feedback | 0.050 | 165/288 (57.3%) | 58/288 (20.1%) | 27/288 (9.4%; [6.5, 13.3]) | 27/58 (46.6%; [34.3, 59.2]) |
| Random feedback | 0.100 | 165/288 (57.3%) | 58/288 (20.1%) | 32/288 (11.1%; [8.0, 15.3]) | 32/58 (55.2%; [42.5, 67.3]) |

### E.5 IFNG Feedback-Granularity Audit

The IFNG audit included five completed five-round trajectories per interface, yielding 20 adjacent-round transitions. Fine feedback returned per-gene/signal results for tested candidates, whereas true feedback returned the aggregate batch and cumulative hit counts used in the main experiment.

Under fine feedback, mean final top-effect recall was 10.96%. Eighteen of 20 transitions stated a feedback-supported adjustment, with 0/20 implemented adjustments and 0/20 outcomes above random expectation. Under true feedback, mean final top-effect recall was 12.17%, with 15/20 supported adjustments, 2/20 implemented adjustments, and 1/20 outcomes above random expectation.

**Table 21:**
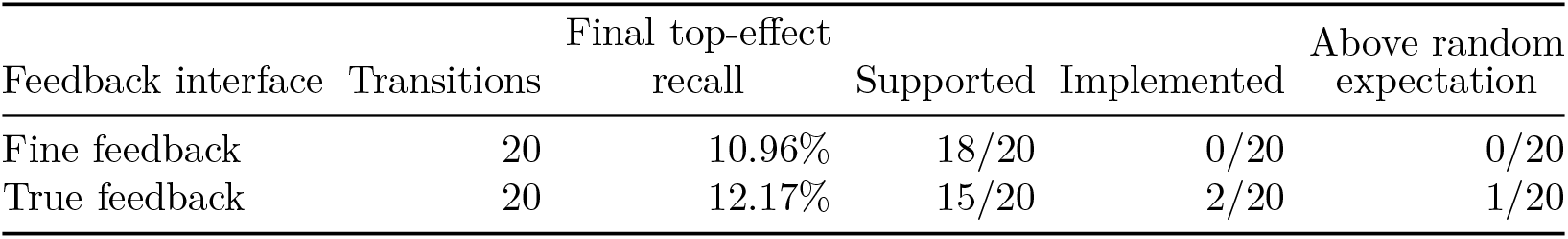
Observable feedback-to-outcome sequences in the IFNG feedback-granularity audit. Fine feedback returns per-gene/signal results for tested candidates, whereas true feedback returns aggregate batch and cumulative hit counts. “Supported” denotes a stated adjustment supported by the displayed feedback; “Implemented” denotes a supported adjustment reflected in the accepted next batch; and “Above random expectation” denotes an implemented adjustment whose next batch contains more benchmark-defined hits than expected under the hypergeometric baseline. Later stages are subsets of earlier stages. Final top-effect recall is the mean across five trajectories per interface.

### E.6 Representative Transition Packets

**Table 22:**
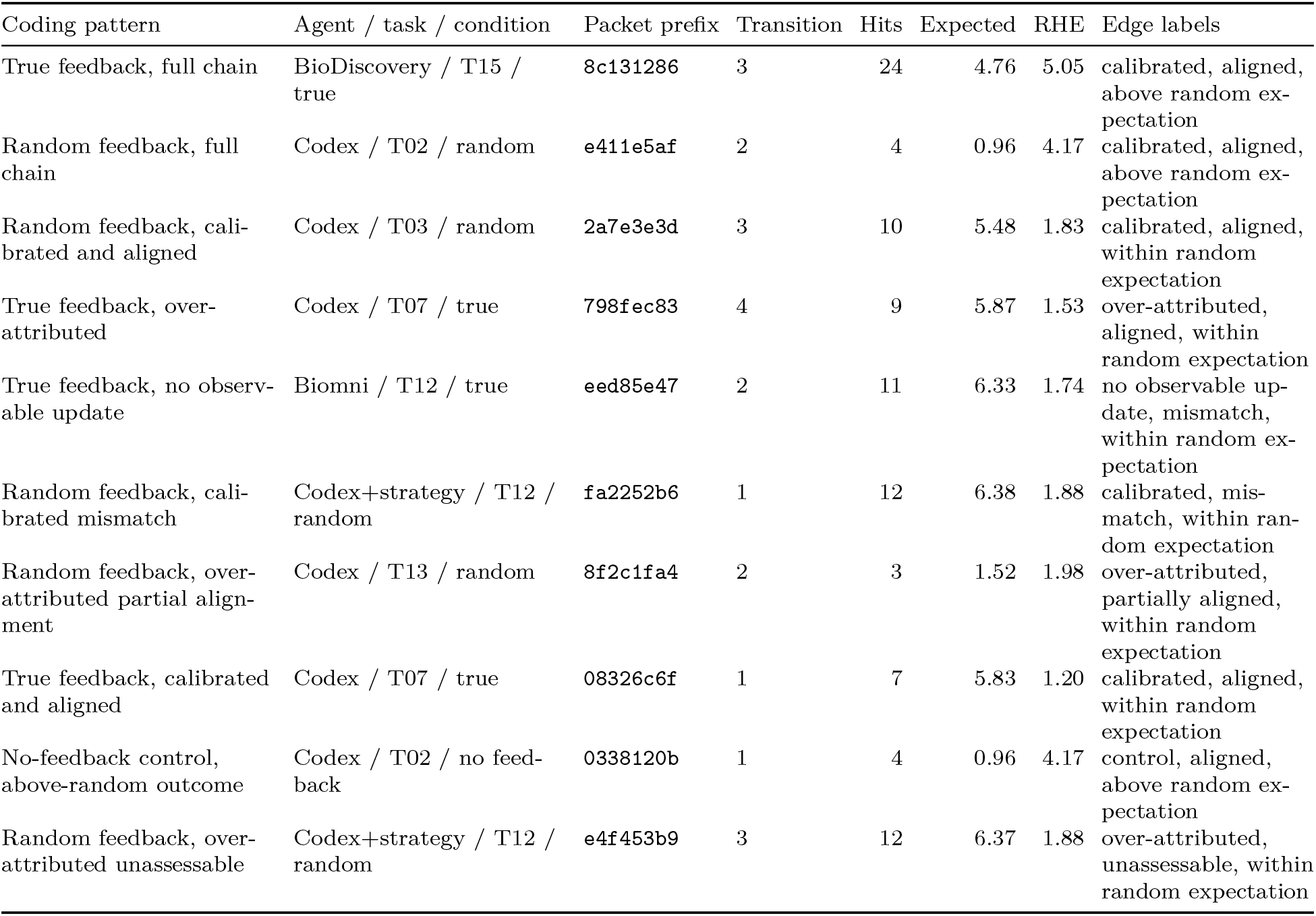
Representative transition packets selected deterministically within each coding pattern. Expected hits are the hypergeometric means from the remaining pool.

## F Prompt Templates

### F.1 Prompt Examples

#### System prompt

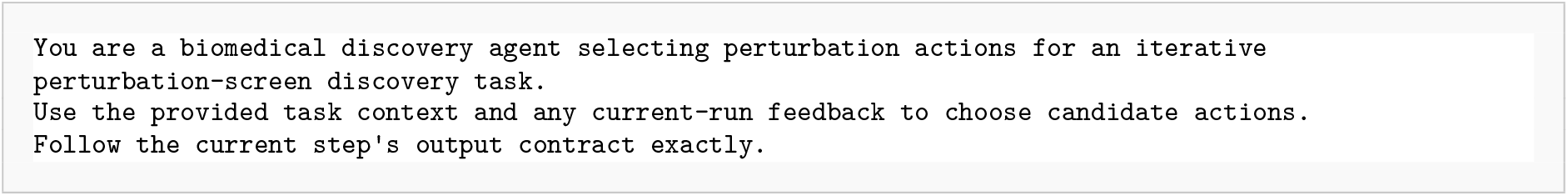

#### Complete round-1 user prompt

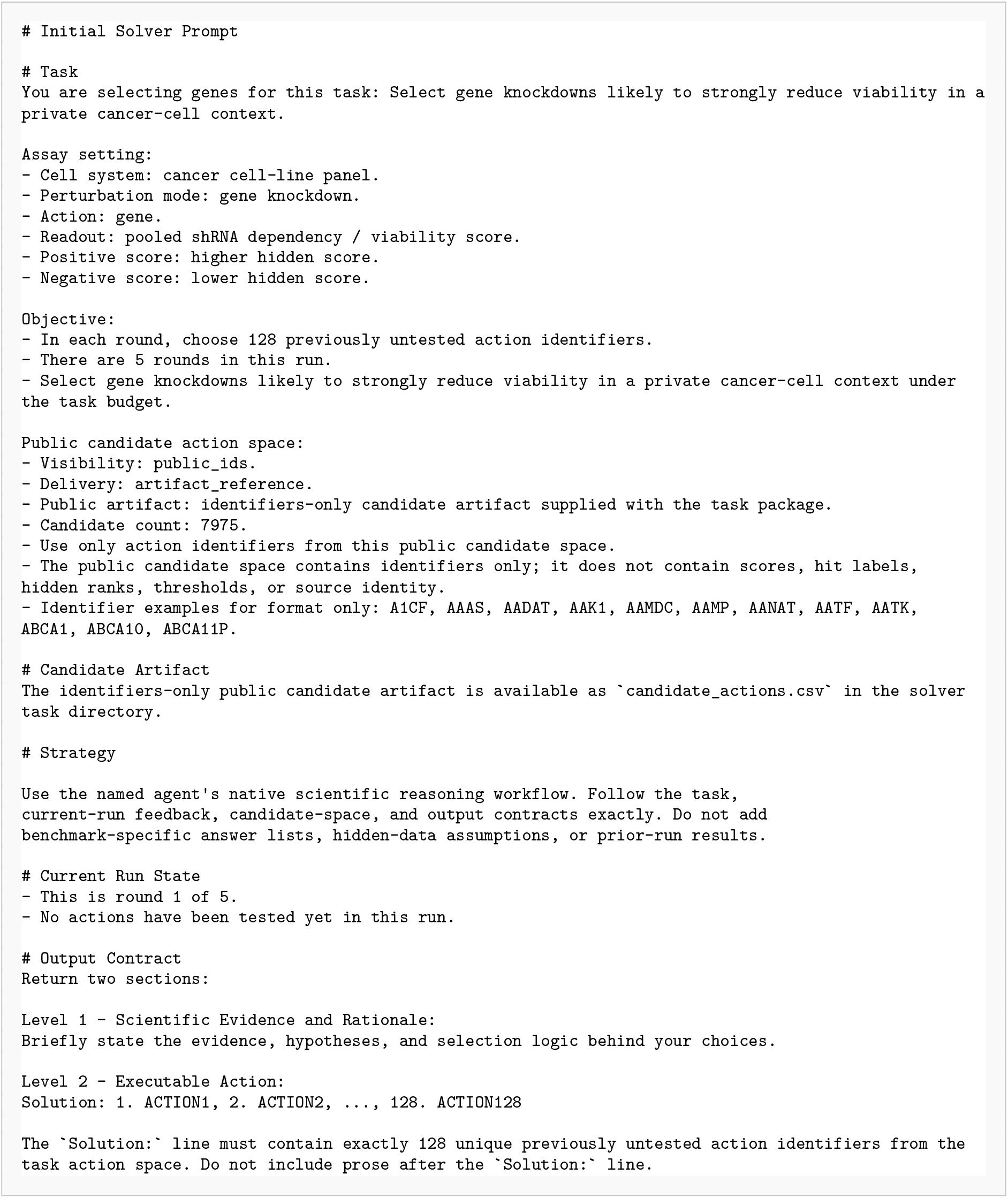

#### Complete strategy prompt for Codex+strategy

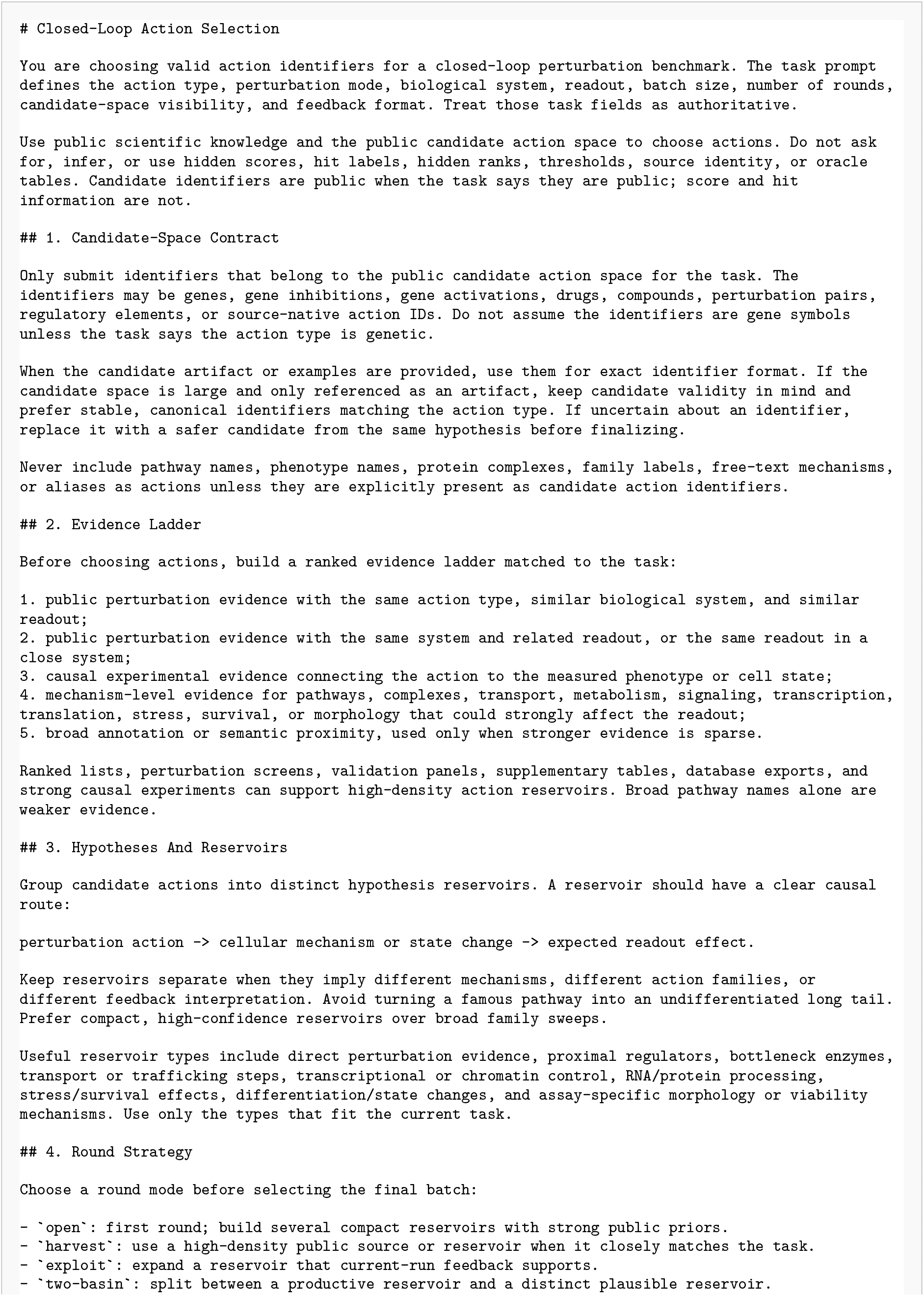
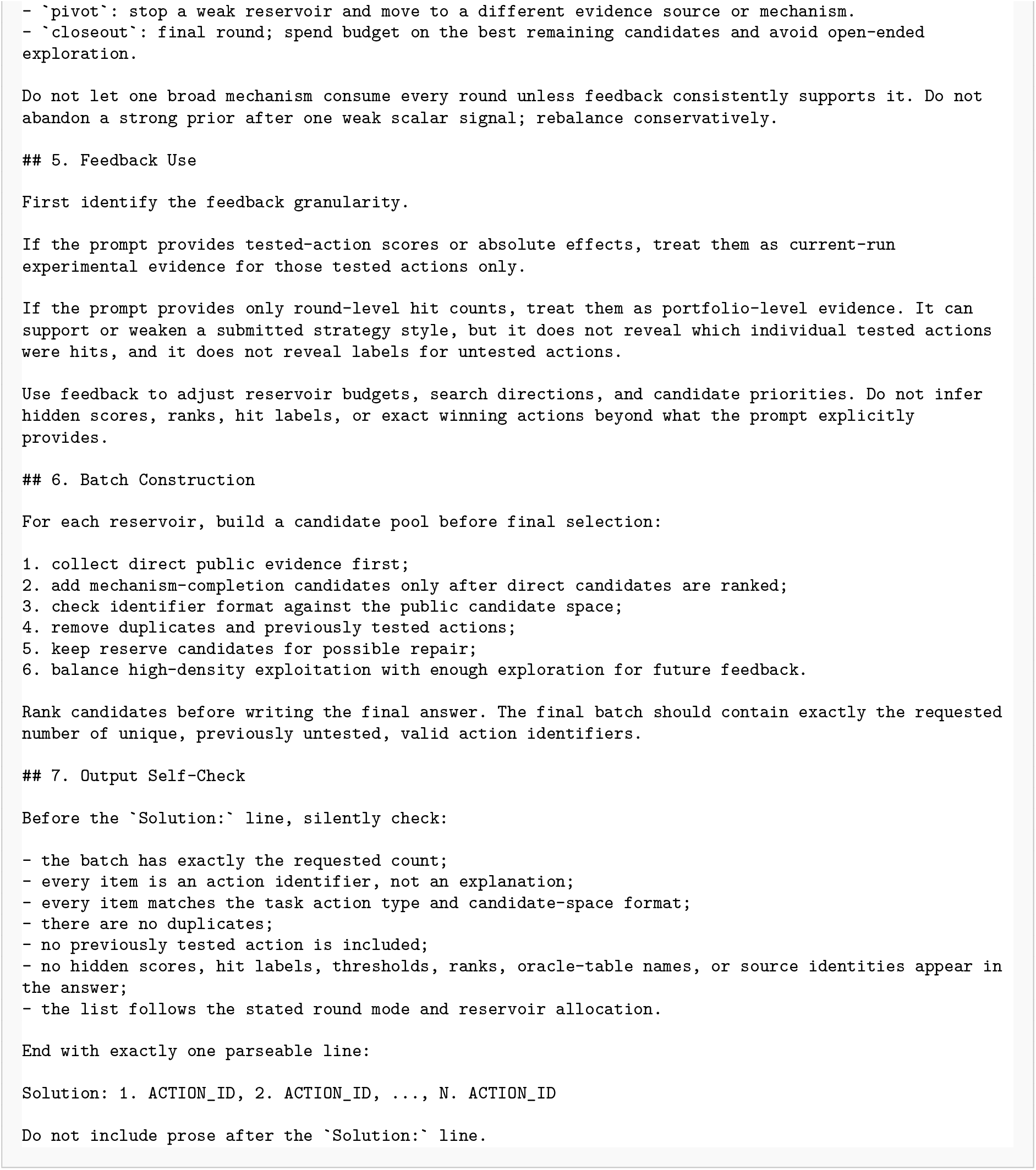

